# Rab2B promotes chaperonin-mediated actin folding and prevents developmental transcription reprogramming and quiescence in trypanosomes

**DOI:** 10.1101/2024.11.27.625582

**Authors:** Feng-Jun Li, Xiaoduo Dong, Yameng Huang, Donghui Hu, Eunyoung Chae, Cynthia Y. He

## Abstract

The Rab family of small GTPases are regulators of intracellular membrane trafficking. Rab2B, a Golgi-resident Rab GTPase, has been shown to function in ER-Golgi transport and autophagosome-lysosome fusion in mammalian cells and *Drosophila*. In *Trypanosoma brucei*, the autophagy-related function of Rab2B was conserved and depletion of Rab2B blocked autophagosome-lysosome fusion. Intriguingly, depletion of Rab2B induced differentiation of *T. brucei* from a proliferative slender form to a quiescent stumpy form, a step crucial for parasite transmission. Chaperonin CCT/TRiC was identified as an effector of Rab2B. Depletion of CCT/TRiC subunits or actin (a known CCT/TRiC substrate), or blocking actin polymerisation with Latrunculin A all triggered differentiation, suggesting a role of Rab2B on CCT/TRiC-mediated actin folding. Using an actin chromobody, we identified a pool of nuclear actin that is crucial for transcription reprogramming and life cycle differentiation in *T. brucei.* The study revealed a function of Rab2B and actin in *T. brucei* transcription reprogramming leading to cell differentiation, and an unexpected role of Rab2B and CCT/TRiC mediated actin folding.

## INTRODUCTION

Hallmarks of cellular differentiation include interplay of various signaling pathways and remodeling of transcriptional and translational activities. Parasitic protozoa such as Plasmodium and trypanosomes also respond to extracellular cues to regulate their proliferation, tropism and development to ensure successful transmission between vectors and hosts^1,2^. However, due to the evolutionary divergence, the coherent regulatory pathway and the connection to tissue tropism and developmental adaption remain poorly understood in most pathogens. Subspecies of *Trypanosoma brucei* are causative agents for African Sleeping Sickness (trypanosomiasis) in humans and nagana in cattle. The life cycle alternates between mammalian hosts and tsetse fly vector, with the parasites taking on different forms and metabolic functions adapted to different environmental niches. Within the complex life cycle, the slender-to-stumpy differentiation step that occurs in mammal blood is of particular interest as a main point of therapeutic intervention^3^.

During mammal infection, *T. brucei* proliferates as slender form in the blood. Once a critical cell density is reached, the slender cells sense an environmental cue known as stumpy induction factor (SIF), and initiate differentiation into a quiescent stumpy form^4^. The stumpy cells are not proliferative, but have remodelled gene expression and metabolism that preadapt the parasite for further development and proliferation in the tsetse fly midgut as procyclic form^4,5^. Recent studies show that oligopeptides produced by secreted trypanosomal oligopeptidases, function as SIF^6^. Oligopeptides are perceived by an orphan GPCR-like protein TbGPR89, consequently driving the slender to stumpy transformation via an unknown mechanism involving protein kinases, protein phosphatase, and RNA-binding proteins RBP7 and ZC3H20^7–9^.

Autophagy, a pathway that clears autophagosome-engulfed macromolecules or organelles through lysosomes, has been found involved in bulk morphological and metabolic changes occur during development and cell differentiation^10^. Autophagy can be triggered upon inhibition of the target of rapamycin (TOR), or activation of AMP-activated protein kinase (AMPK), leading to nucleation, phagophore formation, autophagosome formation/maturation, autophagosome-lysosome fusion, and final degradation. As a continuous and dynamic vesicle transport process, autophagy requires the activity of several Rab small GTPases at different steps. While Rab1, 5, 7, 9A, 11, 23, 32 and 33b are involved in autophagosome formation/biogenesis and maturation^11–17^, Rab2, 7, 33b and 39A are crucial for autophagosome-lysosome fusion and followed autophagosome degradation^18–26^.

Autophagy has also been reported during starvation and life cycle developments in different trypanosomatids species^27–30^. Despite conservation of the core Atg8-Atg7-Atg3 ubiquitin-like conjugation system, only half of known autophagy related genes (ATGs) are found in *T. brucei* genome^31^. Various studies suggest a non-canonical autophagy pathway in *T. brucei* that may be an evolutionary adaptation to its parasitic lifestyle^32^. In this study, we performed an RNAi screening against all 16 Rab proteins in *T. brucei* and identified Rab2B as an autophagy regulator that facilitates autophagosome-lysosome fusion. Surprisingly, depletion of Rab2B in bloodstream form (BSF) trypanosomes led to cell differentiation from the proliferative slender form to the quiescent stumpy form. The differentiation induced by Rab2B depletion was autophagy-independent. We identified the chaperonin complex, also known as CCT(Chaperonin containing TCP-1) or TRiC (T-complex protein Ring Complex), to be a Rab2B effector. Depletion of CCT/TRiC or its substrate actin both induced slender-to-stumpy differentiation, phenocopying Rab2B depletion. These results demonstrated an unexpected role of Rab2B and actin as negative regulators of slender-to-stumpy differentiation in *T. brucei*, likely via CCT/TRiC-mediated actin folding.

## RESULTS

### *T. brucei* Rab2B has a conserved function in autophagosome-lysosome fusion

Previous studies identified 15 distinct Rab GTPase genes (and a truncated form of Rab1A) in *T. brucei* genome^33^. Among them, Rab1A, Rab2B, Rab5 and Rab11 have been shown to play important roles in endocytosis and membrane antigen trafficking/recycling^34–36^. To investigate the role of membrane trafficking in *T. brucei* autophagy, an RNAi-based screening against all Rab GTPases was performed in procyclic cells expressing YFP-tagged ATG8.2 as an autophagosome marker^29^. RNAi of three Rab GTPases, Rab1A, Rab2B and Rab11 were found to have significant effects on starvation-induced autophagy. When cultivated in the medium, basal level autophagy was observed in control cells with ∼0.2 autophagosome/cell. Knockdown (KD) of Rab2B or Rab11 led to an increase to > 2 autophagosomes/cell in the medium and further increase upon starvation (Fig. 1a, b; Supplementary Fig. 1d, e). Rab1A KD in contrary, inhibited starvation-induced autophagosome formation (Supplementary Fig. 1a). Depletion of Rab1A or Rab11 were both lethal (Supplementary Fig. 1a, c). Rab11 KD led to multinucleated cells likely due to cell division defects (Supplementary Fig. 1e). As such for Rab1A and Rab11, we could not rule out the possibility that the autophagy effects observed in these cells were secondary to cytokinesis defects and/or cell death. Rab2B was the only Rab GTPase with a significant effect on autophagy and yet non-essential for cell survival in culture, and it was therefore further investigated in this study.

**Fig. 1:**
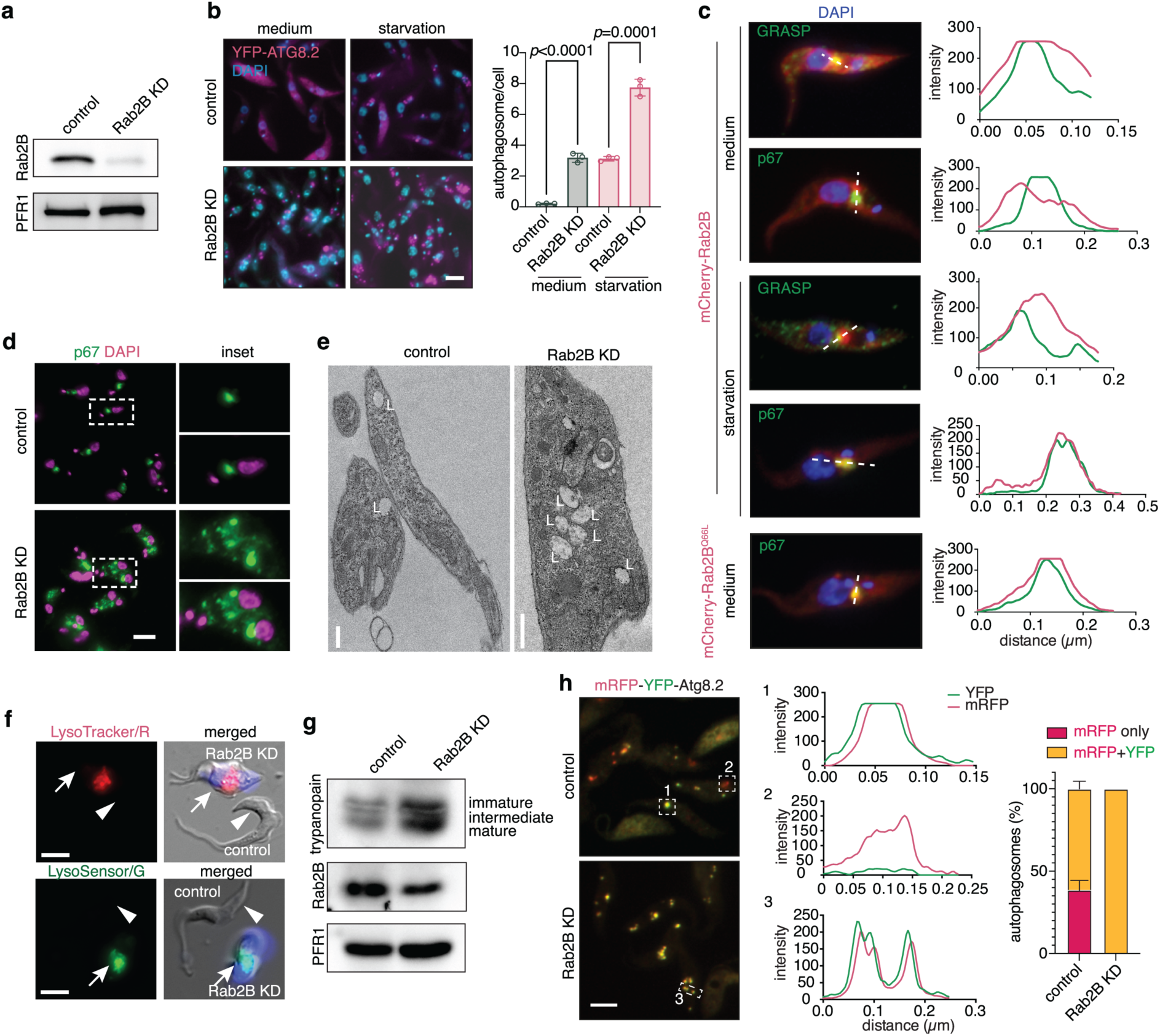
Rab2B is a regulator of autophagy in *T. brucei*. **a)** Rab2B RNAi was induced in PCF cells with 10 µg/ml tetracycline for 48 h and cell lysates were analyzed by SDS-PAGE followed with immunoblotting with a rat polyclonal antibody against *T. brucei* Rab2B. The flagellar protein PFR1 was used as loading control. **b)** RNAi of Rab2B in PCF *T. brucei* led to an increase in autophagosome number under both fed and starvation conditions. YFP-tagged TbATG8.2 was used as autophagosome marker. 48 h post induction of Rab2B RNAi, control (cells cultivated in medium) or cells starved for 2 h in gHBSS were processed for fluorescence microscopy and the autophagosome number was quantitated. Number of autophagosomes/cell was shown as mean ± SD from three independent experiments. More than 200 cells were quantitated in each of the experiment. *p* values were calculated using 2 tailed, unpaired t-test (Prism GraphPad, ver. 10). Scale bar: 5 µm. **c)** The localization of Rab2B in *T. brucei*. Cells expressing mCherry-tagged Rab2B were cultivated in medium or starved for 2 h, fixed and immunolabelled with anti-p67 (for lysosomes) or anti-GRASP (for Golgi). GTP-locked Rab2B^Q66L^ mutant localized to the lysosome even under fed condition. Colocalization of Rab2B with lysosome or Golgi was demonstrated by plot profiles. **d)** Multiple lysosomes were found in BSF cells upon Rab2B RNAi. Anti-p67 antibody was used to stain the lysosomes. Scale bar: 5 µm. **e)** Transmission electron microscopy (TEM) analysis showing multiple lysosomes in Rab2 KD cells compare with control. Scale bars: 200 nm. **f)** Control and Rab2B RNAi cells were stained with LysoTracker Red or LysoSensor Green, the Rab2B RNAi cells were co-stained with 5 µg/ml DAPI. After thoroughly washing with PBSG, control and DAPI-stained Rab2B RNAi cells were then mixed at 1:1 ratio before image acquisition. Scale bars: 5µm. **g)** Tracking the maturation of cathepsin L-like cysteine protease trypanopain by immunoblotting with anti-trypanopain antibody. Three bands were revealed by anti-trypanopain antibody, representing the immature (upper), intermediate (middle) and mature (lower) forms. **h)** Autophagy flux assay. Control or Rab2B KD cells stably expressing mRFP-YFP-TbATG8.2 were starved for 4 h in gHBSS to induce autophagy. YFP and mRFP were monitored in live cells by fluorescence microscopy. The presence of mRFP or YFP signal in autophagosomes was shown by plot profiles and the percentage of autophagosomes containing mRFP only or both mRFP and YFP signals was plotted as mean ± SD of 3 independent experiments. At least 300 autophagosomes were measured in each experiment. Scale bars: 5µm.

Rab2 is present on the Golgi apparatus in mammals and fruit fly^24,37^. In *T. brucei*, N-terminal mCherry-tagged Rab2B colocalized with Golgi protein GRASP^38^, adjacent and distinct to the lysosome marker p67^39^ (Fig. 1c). However, upon starvation, Rab2B co-localized with p67 on lysosomes (Fig. 1c). This Golgi-to-lysosome relocalization is likely due to Rab2B activation during starvation, as the constitutively active Rab2B^Q66L^ mutant localized to the lysosome even under fed conditions (Fig. 1c). In Rab2B-depleted cells, the number and structure of the Golgi appeared unchanged (Supplementary Fig. 1f) while multiple lysosomes, instead of one in control cells, dispersed in the region between the kinetoplast (mitochondrial DNA aggregate) and the nucleus (Fig. 1d, Supplementary Fig. 1g-h). The increasing of lysosome number was further confirmed by transmission electron microscopy (Fig. 1e). These observations are generally consistent with a previous study^36^. Interestingly, the trafficking of lysosomal membrane glycoprotein p67 (Ref.^40^) from Golgi to lysosome appeared to be blocked in Rab2B-KD cells, leading to the accumulation of p67 on cell surface (Supplementary Fig. 1i-k). However, the acidity of lysosomes was even enhanced (Fig. 1f), and the maturation of cathepsin L appeared normal (Fig. 1g).

To determine which autophagy step was affected by Rab2B KD, we performed autophagy flux assays using cells expressing mRFP-YFP-tagged ATG8.2. Quenching of the pH-sensitive YFP fluorescence is used as a reporter for autophagosome fusion with lysosome^41^. In control cells, most of the autophagosomes (>60%) contained both mRFP and YFP fluorescence signals. Approximately 35% of autophagosomes displayed mRFP signal only, indicating fusion with lysosomes (Fig. 1h). In Rab2B-KD cells, all autophagosomes were positive for both mRFP and YFP signals. Given that lysosomes in Rab2B-deficient cells displayed acidic pH (see above), our results support a conserved function of Rab2B as a regulator of autophagy, acting at the autophagosome-lysosome fusion step.

### Rab2B KD induces slender-to-stumpy differentiation in bloodstream forms

Rab2B KD was further investigated in slender, BSF cells engineered for tetracycline- and cumate-inducible expression^42,43^. Here Rab2B KD led to rapid inhibition of cell proliferation (Fig. 2a, b), and accumulation of cells arrested at post-mitotic stage with stumpy-like morphology (Fig. 2c). Compared to control cells, the mitochondria in Rab2B-depleted cells appeared more branched (Fig. 2d), suggesting mitochondrial remodelling, a hallmark of cell differentiation from slender to stumpy form^3^.

**Fig. 2:**
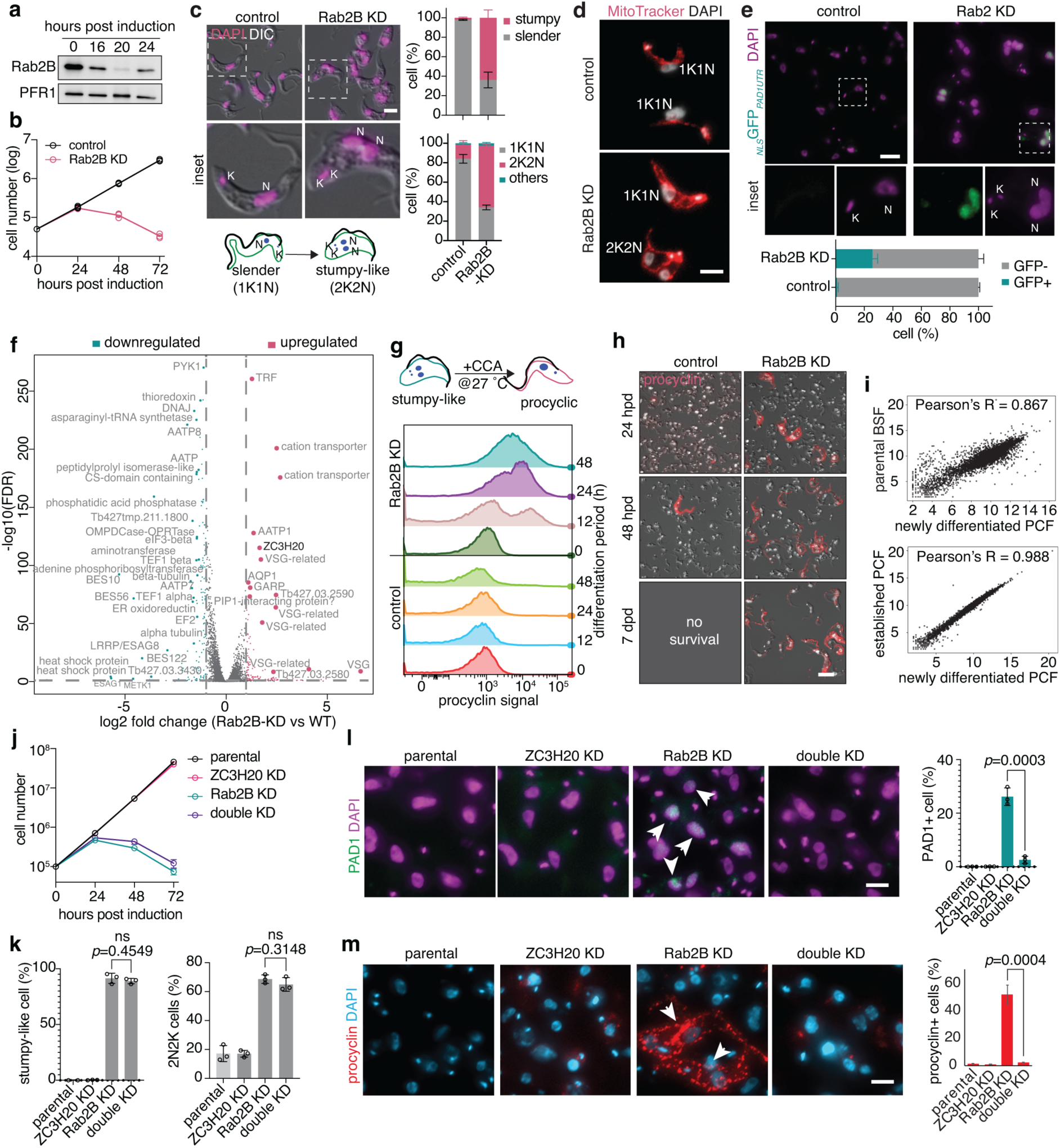
Rab2B depletion triggers slender-to-stumpy differentiation through the SIF pathway. **a)** Rab2B RNAi was induced in BSF cells with 1 µg/ml tetracycline for 16, 20 or 24 h. Cell lysates were fractionated by SDS-PAGE followed by immunoblotting with anti-Rab2B antibody. PFR1 was used as loading control. **b)** Rab2B-depletion reduced BSF cell proliferation in culture. **c)** Rab2B RNAi (20 h) led to changes in the morphology and cell cycle (tracked by DAPI staining) change upon Rab2B RNAi (20 h). The quantification was shown as mean ± SD of 3 independent experiments. At least 200 cells were quantified for each experiment. N: nucleus; K: kinetoplast. Scale bar: 5 µm. **d)** Changes in mitochondria morphology in Rab2B KD cells as revealed by MitoTracker Red staining. N: nucleus; K: kinetoplast. Scale bar: 5 µm. **e)** The expression of stumpy marker PAD1 in control and Rab2B-depleted cells. Cells stably expressing PAD1 3’ UTR regulated eGFP fused to a nuclear localization sequence were induced for Rab2B RNAi or not. The quantification of eGFP positive cells in control and Rab2B KD cells was shown as mean ± SD of 3 independent experiments. At least 200 cells were quantified for each experiment. **f)** Volcano plot displaying differentially expressed genes in Rab2B KD cells compared with the uninduced control cells. The cyan and pink dots represent genes with expression levels downregulated or upregulated, respectively, with fold change>2. The grey dots represent transcripts with fold change<2. **g)** The Rab2B KD-induced stumpy cells can be efficiently differentiated into procyclic forms *in vitro*. Control and Rab2B KD cells (20 h post RNAi induction) were resuspended in Cunningham’s medium containing 6 mM CCA to a density of 3–5×10^6^/ml, and cultivated at 27 °C. At each indicated time point, the cells were collect, stained with EP1 antibody that targets the surface procyclin and analysed by flow cytometry. **h)** Immunofluorescence microscopy shows the expression of cell surface antigen procyclin (EP1) during the in vitro differentiation experiment as described in (G). Scale bar: 5 µm. **i)** Comparison of transcriptomes of new differentiated PCF cells from the experiments described in (G) and (H) to parental BSF cells and lab adapted PCF YTat1.1 cells. *R* indicated the Pearson’s correlation coefficient. **j)** Growth assays of Rab2B RNAi, ZC3H20 RNAi and the double RNAi cells. ZC3H20 knockdown could not rescue the lethal phenotype caused by Rab2B RNAi. **k)** The quantification of stumpy-like cells and cells with cell cycle arrested at 2N2K phase upon individual or double RNAi of Rab2B and ZC3H20. The data were displeayed as mean ± SD for 3 independent experiments. At least 200 cells were quantified in each experiment. **l)** The expression of PAD1 in control and cells induced for Rab2B and ZC3H20 RNAi, individually or together. The quantification of PAD1-positive cells was displayed as mean ± SD for 3 independent experiments. At least 200 cells were quantified in each experiment. Scale bar: 5 µm. **m)** EP1 expression during *in vitro* differentiation was monitored by fluorescent microscopy. The quantification of procyclin positive cells 48 h post *in vitro* differentiation following individual or double RNAi was shown as mean ± SD for 3 independent experiments. At least 200 cells were quantified in each experiment. Scale bar: 5 µm.

Protein Associated with Differentiation 1 (PAD1) is widely used as a molecular marker for the stumpy form cells^44–46^. Using a reporter vector that expresses nuclear targeted GFP (_NLS_GFP) under the control of PAD1 3’UTR^47^, ∼20% of Rab2B KD cells showed GFP expression in the nucleus at 22 h post RNAi-induction (Fig. 2e). RNA-seq analyses of Rab2B-KD cells showed a transcriptomic profile highly similar to that published for stumpy forms^48^. This included upregulation of many VSG and expression site associated genes, aquaporins, two cation transporters and RNA binding protein ZC3H20; and downregulation of transcripts encoding flagellar proteins, histones, and alpha/beta-tubulin (Fig. 2f). These results suggested that Rab2B depletion triggered slender-to-stumpy differentiation in monomorphic BSF trypanosomes, which cannot sense and/or respond to SIF and therefore lose the ability to differentiate spontaneously^49^.

Importantly, Rab2B KD-induced stumpy cells were competent for further differentiation to procyclic form. Upon the addition of 6 mM citrate/cis-aconitate (CCA) and temperature shift to 27 °C^49^, procyclin-positive cells were occasionally observed in control cells 24 or 48 hrs post differentiation, but no viable cells could be found after 7 days (Fig. 2g, h). Rab2B KD cells however, exhibited increasing fraction of procyclin-positive cells from 18.6 ± 1.98% at 24 hrs, to 49.8 ± 4.52% at 48 hrs (Fig. 2g, h). After 7 days, the cells proliferated robustly in culture and all cells are procyclin-positive (Fig. 2g, h). These differentiated procyclic cells can be cultured continuously as a stable cell line. Transcriptomic profile of these newly differentiated procyclic cells was near identical to that published previously for lab-adapted procyclic cells^48^ (*R*=0.998; Fig. 2i, lower panel) but more different to the parental slender cells (*R*=0.867; Fig. 2i, upper panel).

Comparing the transcriptomes of the newly differentiated procyclic cells with those of the parental slender cells, we identified approximately 400 or 800 genes that are up- or down-regulated, respectively, with a fold change >2 and *p* value of < 0.05 (Supplementary Fig. 2a). Distinct RNA expression pattern was also apparent in PCA analyses (Supplementary Fig. 2b). Among the downregulated genes, about 30% were VSGs and expression site related genes (ESAGs) (Supplementary Fig. 2c). In upregulated genes, procyclins and the genes involved in mitochondria metabolism or respiration were enriched (Supplementary Fig. 2c). GO analysis further indicated upregulation of TCA cycle- and cellular respiration-related genes and downregulation of genes involved in glucose metabolism or glycolysis (Supplementary Fig. 2d), all consistent with the metabolic profile of a procyclic form. Together, these results demonstrated that Rab2B-depletion triggered monomorphic slender cell differentiation into stumpy form, which can further differentiate into viable procyclic cells *in vitro*.

### Rab2B KD-induced differentiation requires ZC3H20

In the mammal blood, slender-to-stumpy differentiation takes place via a quorum-sensing mechanism in response to stumpy inducing factor (SIF). The SIF pathway involves the activities of several kinases/phosphatases and two RNA-binding proteins, RBP7 and ZC3H20^49^. The expression of ZC3H20 is deemed as a key event marking the commitment of cell differentiation to stumpy form^9^. A significant increase in the transcription level of ZC3H20 was observed in Rab2B-KD cells (Fig. 2f), raising a possibility that Rab2B KD-induced differentiation was via the SIF pathway. To investigate the function of ZC3H20 in Rab2B KD-induced differentiation, cells with double KD of both Rab2B and ZC3H20 were generated. ZC3H20 depletion was not able to rescue the lethal phenotype caused by Rab2B depletion (Fig. 2j). The cells were morphologically stumpy and arrested at post-mitotic stage (Fig. 2k). However, ZC3H20 depletion blocked the expression of the stumpy marker PAD1 (Fig. 2l). These PAD1-negative, morphologically stumpy cells were not committed and could not further differentiate into procyclic cells (Fig. 2m). These findings suggest that Rab2B depletion may induce the slender-to-stumpy differentiation via the SIF pathway.

### Rab2B KD-induced differentiation is autophagy-independent

Autophagy has been shown to drive cell differentiation and metabolic remodelling in many organisms including *T. brucei*^32^. However, as Rab2B depletion inhibited autophagy flux (c.f. Fig. 1h), we reasoned that Rab2B KD-induced slender-to-stumpy differentiation does not require functional autophagy. To determine if cell differentiation can be triggered by autophagy deficiency, we performed RNAi on ATG3, an autophagy-related gene that is required for autophagosome formation^29^. Depletion of ATG3 did not induce any morphological change or PAD1 expression (Supplementary Fig. 3). Together, these results demonstrated that Rab2B KD-induced slender-to-stumpy differentiation is independent of autophagy.

### Chaperonin CCT/TRiC is a Rab2B effector

To further investigate the mechanisms of Rab2B in differentiation, we performed co-immunoprecipitation assays on cells stably expressing YFP-tagged Rab2B^Q66L^ (active form) and Rab2B^S21N^ (inactive form) mutants. Cells expressing YFP-only was used as a negative control. Several protein bands were found enriched in Rab2B^Q66L^ samples (arrows, Fig. 3a). Mass spectrometry identified seven of eight known chaperonin CCT/TRiC subunits, a heat shock protein HSP60, two polyadenylate-binding proteins (PABP1 and PABP2), and one glycosomal transporter GAT2 (Fig. 3b). An RNAi screen was performed to investigate their potential effects on cell growth and differentiation. Among the above proteins, only CCT3 and CCT6, both subunits of the CCT/TRiC complex, triggered the expression of PAD1 upon RNAi (Fig. 3c, d). The association of chaperonin with Rab2B in a GTP-dependent manner was further confirmed by Co-IP (Fig. 3e). These results identified CCT/TRiC as an effector of Rab2B.

**Fig. 3.**
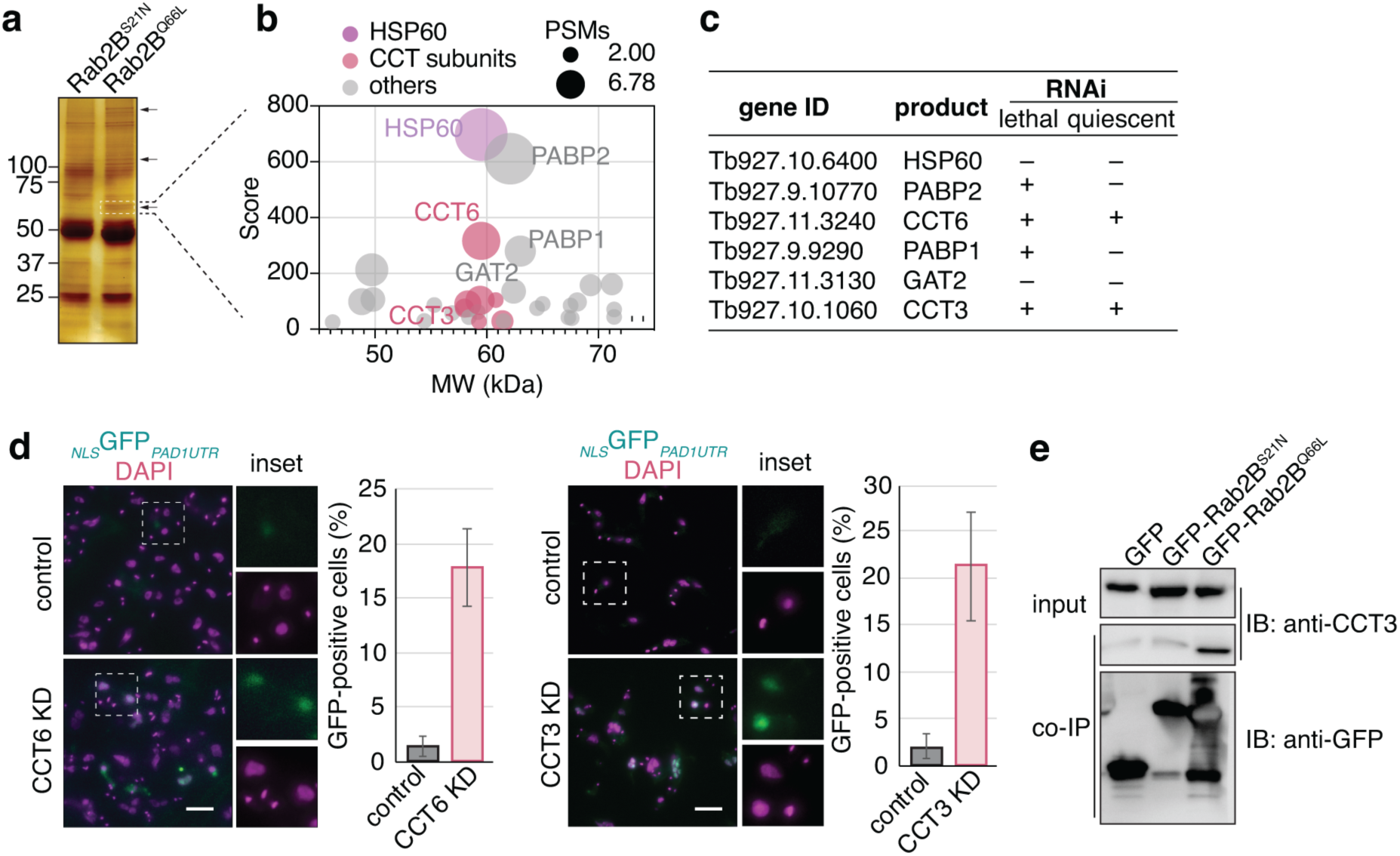
Identification of chaperonin (CCT/TRiC) as a Rab2B effector. **a)** Cleared lysates of YFP-Rab2B^S21N^ and YFP-Rab2B^Q66L^-expressing cells were incubated with magnetic beads conjugated to GFP nanobody (GFP-Trap, ChromoTek). Co-IP products were analyzed by SDS-PAGE followed with silver staining. Several bands were enriched in YFP-Rab2B^Q66L^ but not in YFP-Rab2B^S21N^ (arrows). Regions containing these bands were sliced out and analyzed by mass spectrometry. **b)** Proteomic analyses of one of the regions (marked by dotted lines) identified PABP1, PABP2, and several components of type I (HSP60) and type II (CCT/TRiC) chaperonin as putative Rab2B effectors. **c)** Summary of RNAi analyses of the putative Rab2B effectors. Only CCT3 and CCT6 depletion leads to slender-to-stumpy differentiation. **d)** Depletion of CCT3 or CCT6 triggers PAD1 expression and cell differentiation to stumpy forms. Quantification was shown as mean ± SD for 3 independent experiments. At least 200 cells were quantified in each experiment. Scale bar: 5 µm. **e)** Rab2B interacts with CCT/TRiC in a GTP-dependent manner. Cleared lysates of YFP-Rab2B^S21N^ and YFP-Rab2B^Q66L^-expressing cells were incubated with magnetic beads conjugated to GFP nanobody (GFP-Trap, ChromoTek). Cells expressing GFP only were used as a negative control. Co-IP products were analyzed by SDS-PAGE followed with immunoblots with an anti-CCT3 antibody. GFP and GFP-tagged Rab2B baits were detected by anti-GFP antibody.

### Chaperonin CCT/TRiC-mediated actin folding prevents the slender-to stumpy differentiation

Chaperonin is a protein complex involved in the folding of certain proteins and is highly conserved from bacteria (type I chaperonin GroEL) to humans (type I chaperonin HSP60-HSP10 and type II chaperonin CCT/TRiC/TCP-1)^50^. In *T. brucei*, the CCT/TRiC complex contains eight subunits, which are highly conserved in primary sequence (>50% identity) compared to orthologues found in other eukaryotes (Supplementary Fig. 4a, b). All *T. brucei* CCT/TRiC subunits were predicted to contain apical, intermediate, and equatorial domains (Supplementary Fig. 4c), similar to their mammal orthologues.

While approximately 10% of total proteins show interaction with CCT/TRiC, only few of them have been confirmed as folding substrates^50^. These include cytoskeletal proteins actin and tubulin, and a family of proteins containing 7-bladed WD-domains, including cell cycle regulators CDC20 and Cdh1, G-protein subunit Gβ and mTOR complex components mLST8 and Raptor^50^. Actin and tubulin (α and β) are well conserved in *T. brucei*. Three homologs of 7-bladed WD-domain-containing proteins Raptor, LST8, and CDC20 were also found in *T. brucei* genome (Fig. 4a). Of these five putative CCT/TRiC substrates, only actin depletion showed differentiation phenotypes similar to that observed for Rab2B-RNAi cells (Fig. 4b). In *T. brucei,* actin filament is likely rare and transient, as filament actin has not be observed using phalloidin or other actin visualization methods^51^. To evaluate the effect of CCT/TRiC on actin in *T. brucei*, F-actin to G-actin ratio was examined in control and cells with RNAi depletion of CCT3 or CCT6. Assuming that unfolded actin cannot be polymerized into actin filaments, reduced F-actin/G-actin ratio would suggest chaperonin dysfunction, which was indeed observed in CCT3- and CCT6-depleted cells (Fig. 4c). The cytoskeleton-associated actin was decreased to about 30-40% of control in CCT3- or CCT6-depleted cells (Fig. 4d).

**Fig. 4:**
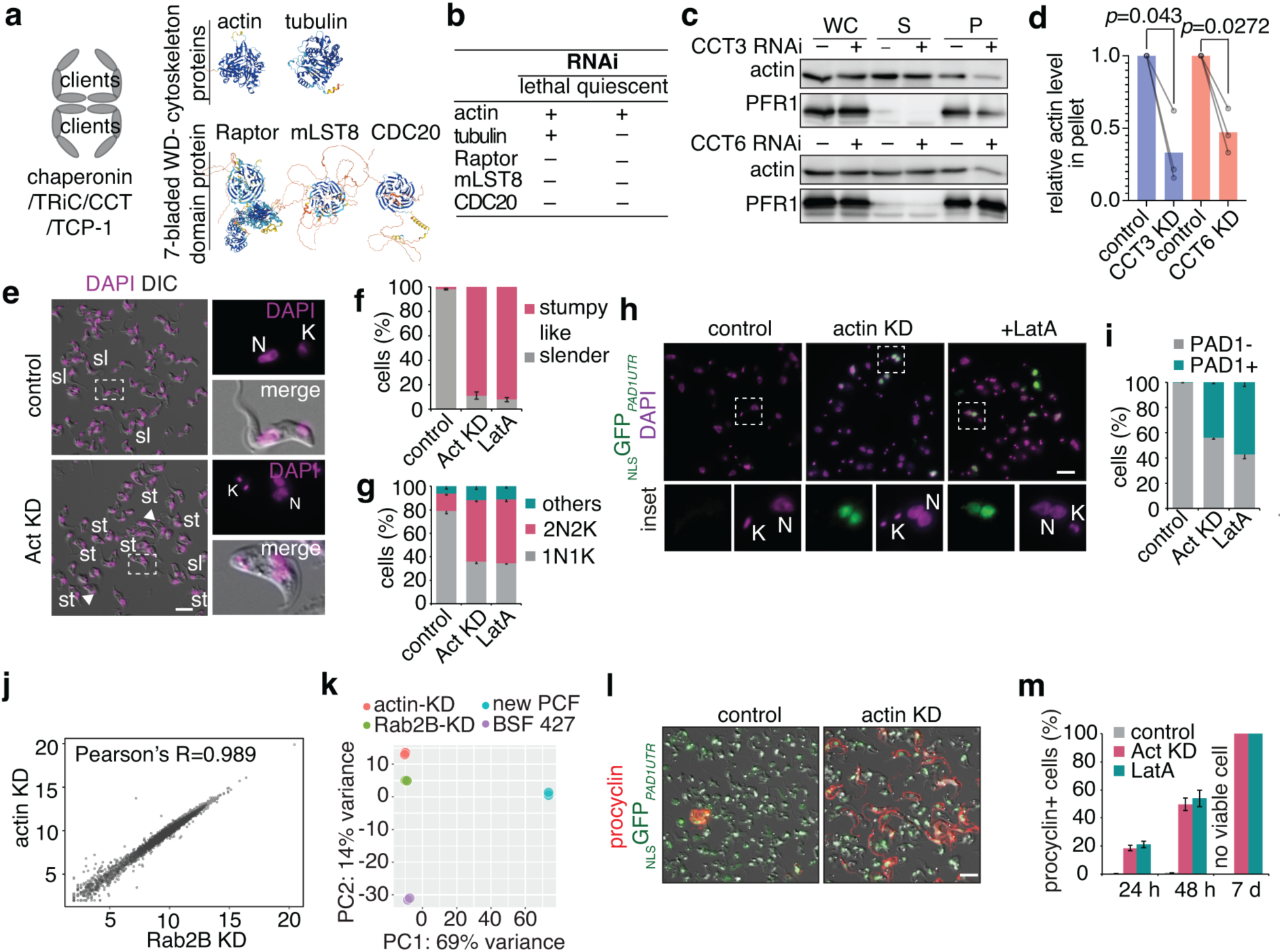
Depletion of actin triggers slender-to-stumpy differentiation. **a)** The orthologues of two classes of well-characterized CCT/TRiC substrates: cytoskeleton proteins actin and tubulin, and 7-bladded WD-domain containing proteins in *T. brucei*. **b)** Summary of the RNAi phenotypes of the potential CCT/TRiC substrates shown in (A). Both actin and tubulin are essential for survival of trypanosomes in culture, but only actin KD induced slender-to-stumpy differentiation. **c)** CCT/TRiC depletion reduced cytoskeleton-associated actin. Control or CCT3/CCT6 RNAi cells were treated with PBS containing 1% NP-40 for 30 min. The soluble (S) and insoluble cytoskeletal (P) fractions were separated by centrifugation and the actin levels in each fraction was detected by an anti-actin antibody. RNAi did not affect total actin protein level in the whole cell lysates (WC). Flagellar cytoskeleton protein PFR1 was used as fractionation and loading control. **d)** The amount of cytoskeleton-associated actin was quantified by calculating the ratio of insoluble actin in CCT3- or CCT6-depleted cells and that in control cells. Results from three independent experiments were shown for each cell line. *p* values were calculated by 2 tailed, paired t-test using Prism GraphPad (ver. 10). Scale bar: 5 µm. **e)** Actin RNAi induced stumpy-like cells and cell cycle arrest, similar to that observed in Rab2B RNAi cells. sl: slender; st: stumpy-like; N: nucleus; K: kinetoplast. Scale bar: 5 µm. **f)** Quantification of stumpy-like cells upon actin-RNAi or treatment with actin polymerization inhibitor LatA. The results were shown as mean ± SD for 3 independent experiments. At least 200 cells were quantified in each experiment. **g)** Quantification of cells in different cell cycle stages in control, actin RNAi or cells treated with LatA. The results were shown as mean ± SD for 3 independent experiments. At least 200 cells were quantified in each experiment. **h)** Actin knockdown or LatA treatment both triggers the expression of stumpy marker PAD1. Scale bar: 5 µm. **i)** Quantification of PAD1 positive cells in actin RNAi (24 h post induction) or LatA treated (4 h) cells shown as mean ± SD for 3 independent experiments. At least 200 cells were quantified in each experiment. **j)** Comparison of the relative expression level of genes (dots) in Rab2B KD and actin KD cells. *R*: the Pearson’s correlation coefficient. **k)** PCA analyses showing little transcription variation between the Rab2B KD and actin KD cells, when compared to the parental BSF cells and to the PCF cells. **l)** The stumpy cells induced by actin-depletion or LatA-treatment differentiate further to procyclic form. Wildtype, actin KD (24 h post induction) or LatA-treated (4 h) cells were washed and resuspended in Cunningham’s medium to a density of 3–5×10^6^ cells/ml. After the addition of 6 mM CCA, cells were incubated at 27 °C for 24 h, 48 h, and 7 days. The expression of procyclin, a PCF marker, was monitored using anti-EP1 antibody. Image shown is representative of actin KD cells at 48 h post differentiation. Scale bar: 5 μm. **m)** The quantitation of EP1-positive cells of the experiments described in (L). Error bar represent the SD of 3 independent experiments. At least 200 cells were quantified in each experiment.

Actin depletion phenocopied Rab2B knockdown. Within 24 hours of actin depletion, approximately 85% of cells transformed from long slender to short stumpy-like form (Fig. 4e, f), ∼40% of cells arrested at the post-mitotic phase with 2 kinetoplasts and 2 nuclei (2K2N, Fig. 4e, g) and ∼50% of cells displaying PAD1 expression (Fig. 4h, i). The similarity between actin KD- and Rab2B KD-induced cellular changes was further demonstrated by their near identical transcriptomic profiles (Pearson’s Corelation factor *R* =0.989) (Fig. 4j) and PCA analysis of their transcriptome variations (Fig. 4k). Upregulated gene transcripts in both Rab2B- and actin-depleted cells also shared similarly enriched GO terms (Supplementary Fig. 5). Furthermore, treatment with Latrunculin A (LatA), a specific actin polymerization inhibitor, also triggered PAD1 expression (Fig. 4g, h), suggesting that actin filament formation is crucial to prevent the differentiation. The stumpy cells derived from actin KD and LatA-treatment could both be further differentiated into procyclic forms (Fig. 4l, m).

CCT/TRiC-mediated folding of actin plays a crucial role in autophagosome clearance/degradation in mammalian cells^52^. To test whether the autophagy phenotype observed in Rab2B KD cells was due to deficiency in CCT/TRiC-mediated actin folding, autophagy activity was examined in CCT6-RNAi or LatA-treated cells. Neither treatments had detectable effects on autophagosome numbers (Supplementary Fig. 6), suggesting that Rab2B function on autophagy was independent of actin in *T. brucei*. Our studies thus suggest two independent functions of Rab2B in *T. brucei*, one on autophagy that is independent of actin; the other on differentiation that is via actin and independent of autophagy.

### Rab2B regulates actin organization in *T. brucei* as well as in mammalian cells

Our results so far suggested a function of Rab2B on actin, possibly via Rab2B interaction with CCT/TRiC. Supporting this, the cytoskeleton-associated actin decreased significantly in Rab2B-depleted cells (Fig. 5a, b), similar to that observed in CCT/TRiC mutants (c.f. Fig. 4c, d). As both Rab2B and actin are highly conserved from trypanosomes to mammals, the function of HsRab2A and HsRab2B, two mammal orthologues to Rab2B, on actin organization were evaluated in HeLa cells. Both HsRab2A and HsRab2B were present on the Golgi apparatus, colocalizing with Golgi matrix marker GM130 (Supplementary Fig. 7a). HsRab2B was additionally present in the nucleus. Expression of the inactive S20N mutants dispersed the Golgi structure, while the constitutively active Q65L mutants maintained their Golgi presence but appeared to compact the structure (Supplementary Fig. 7a). This is consistent with HsRab2A and HsRab2B having important functions in Golgi maintenance, and most of their effectors are Golgi-resident proteins^53^. Neither HsRab2A nor HsRab2B, in the active forms, localized to the lysosomes. This is different to the lysosome/late endosome localization of GTP-Rab2B in *T. brucei* (c.f. Fig. 1c) or *Drosophila* S2 cells^53^.

**Fig. 5:**
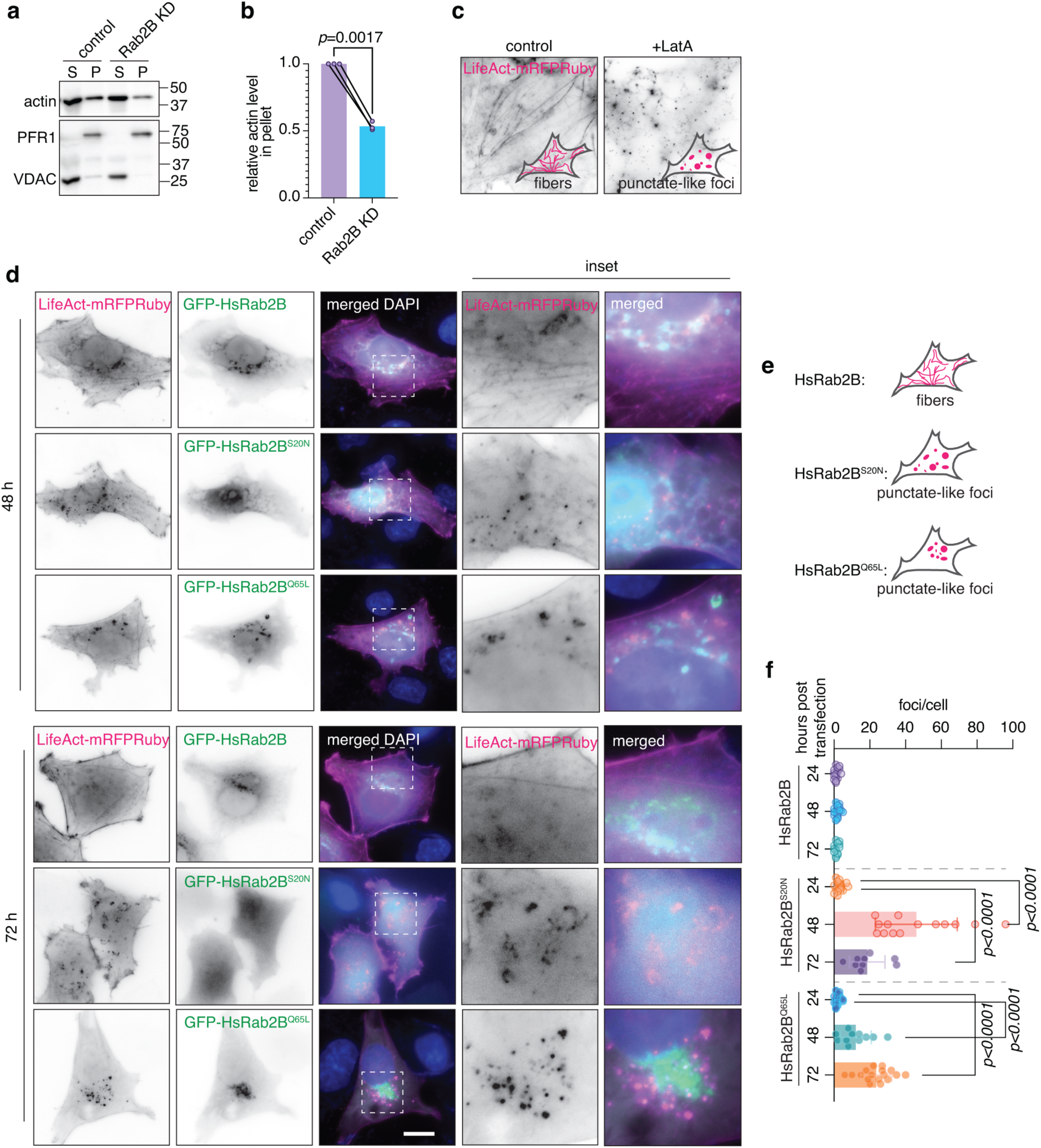
Rab2B regulates actin organization. **a)** The effect of Rab2B KD on actin organization in BSF *T. brucei*. Control and Rab2B RNAi cells were extracted with PBS containing 1% NP-40 for 30 min at room temperature. Whole cell lysate (WC), soluble (S) and insoluble (P) fractions were analyzed by immunoblots and the actin level in each fraction was detected by anti-actin antibody. Flagellar cytoskeleton protein PFR1 and mitochondria membrane protein VDAC were used as controls for loading and detergent extraction. **b)** The effect of Rab2B KD on cytoskeleton-associated actin was quantified by calculating the ratio of insoluble actin in Rab2B-depleted cells and that in control cells. Results from three independent experiments were shown. *p* values were calculated by 2 tailed, paired t-test using Prism GraphPad (ver. 10). **c)** LatA (0.2 µM) treatment leads to changes in actin organization in HeLa cells. Actin organization was tracked by ectopic expression of mRFPRuby-tagged LifeAct in control or LatA-treated (3 h) HeLa cells. **d)** HsRab2B regulates actin organization in HeLa cells. Cells were co-transfected with LifeAct-mRFPRuby and GFP-HsRab2B, wild type, GDP-locked (S20N) or GTP-locked (Q65L). Representative images of cells at 24, 48 or 72 hours post transfection are shown. Scale bar: 10 µm. **e)** Schematic diagrams of actin organization in HeLa cells expressing HsRab2B variants. **f)** Quantification of the number of actin punctate foci in HeLa cells cells at 24, 48 and 72 h post transfection of HsRab2B variants. Each dot represented one cell measured. *p* values were calculated by 2 tailed, unpaired t-test using Prism GraphPad (ver. 10).

Actin filaments was monitored in cells co-expressing GFP-tagged HsRab2A/B mutants and mRFPRuby-tagged LifeAct. Upon LatA treatment, LifeAct-labelled actin fibre was dispersed into punctate-like foci (Fig. 5c), as reported in previous studies.^54^ The actin punctate-like foci were also observed in HsRab2B^S20N^ or HsRab2B^Q65L^ expressing cells, more prominent at 48 or 72 h post transfection (Fig. 5d-f). In comparison, in cells expressing HsRab2A^S20N^ or HsRab2A^Q65L^, actin filament appeared normal up to 72 h post transfection (Supplementary Fig. 7b). Together these results supported a conserved function of Rab2B on actin organization, from trypanosomes to humans.

### Nuclear actin-dependent RNAP2 transcription prevents quiescence/ differentiation

Actin is known for its diverse roles in cellular processes such as cell movement, structural support and endocytosis. The presence of a nuclear pool of actin and their functions in chromatin remodelling and differentiation-related gene regulation is also reported in recent years^55–58^. As life cycle differentiation in *T. brucei* involves significant transcriptional reprogramming^49^, we hypothesized that trypanosome actin may have a role in transcription reprogramming, consequently affecting cell differentiation.

To examine the presence of nuclear actin, recombinant actin nanobody (nAC) fused to TagGFP2 was expressed using a cumate-inducible system in *T. brucei* (Fig. 6a). The ability of the nAC to bind *T. brucei* actin was confirmed by pulldown assays using GST-nAC (Fig. 6b). When expressed in the BSF cells, nAC-TagGFP2 signal was found throughout the cell, with an enrichment in the lysosomes similar to that recently reported^59^. In more than 60% of cells, additional nAC-TagGFP2 enrichment in nuclear foci that appeared to be the nucleolus was also observed (Fig. 6c). Similar nucleolus enrichment was also observed when nAC-TagGFP2 was fused with a nuclear localization sequence (NLS-nAC-TagGFP2) (Fig. 6d). Both lysosome and nucleolus enrichment was abolished in actin-depleted cell (Fig. 6c, d). Together, our results supported the presence of actin in *T. brucei* nucleus. To study the function of nuclear actin, we modified the recently developed targeted protein degradation system (deGradFP)^60^ by replacing the GFP nAb sequence with nAC coding sequence. Targeted degradation of total actin (using degrAdCT) or nuclear actin (using NLS-degrAdCT) both triggered slender-to-stumpy differentiation (Fig. 6e, f), further attesting the role of actin, particularly nuclear actin in *T. brucei* differentiation.

**Fig. 6:**
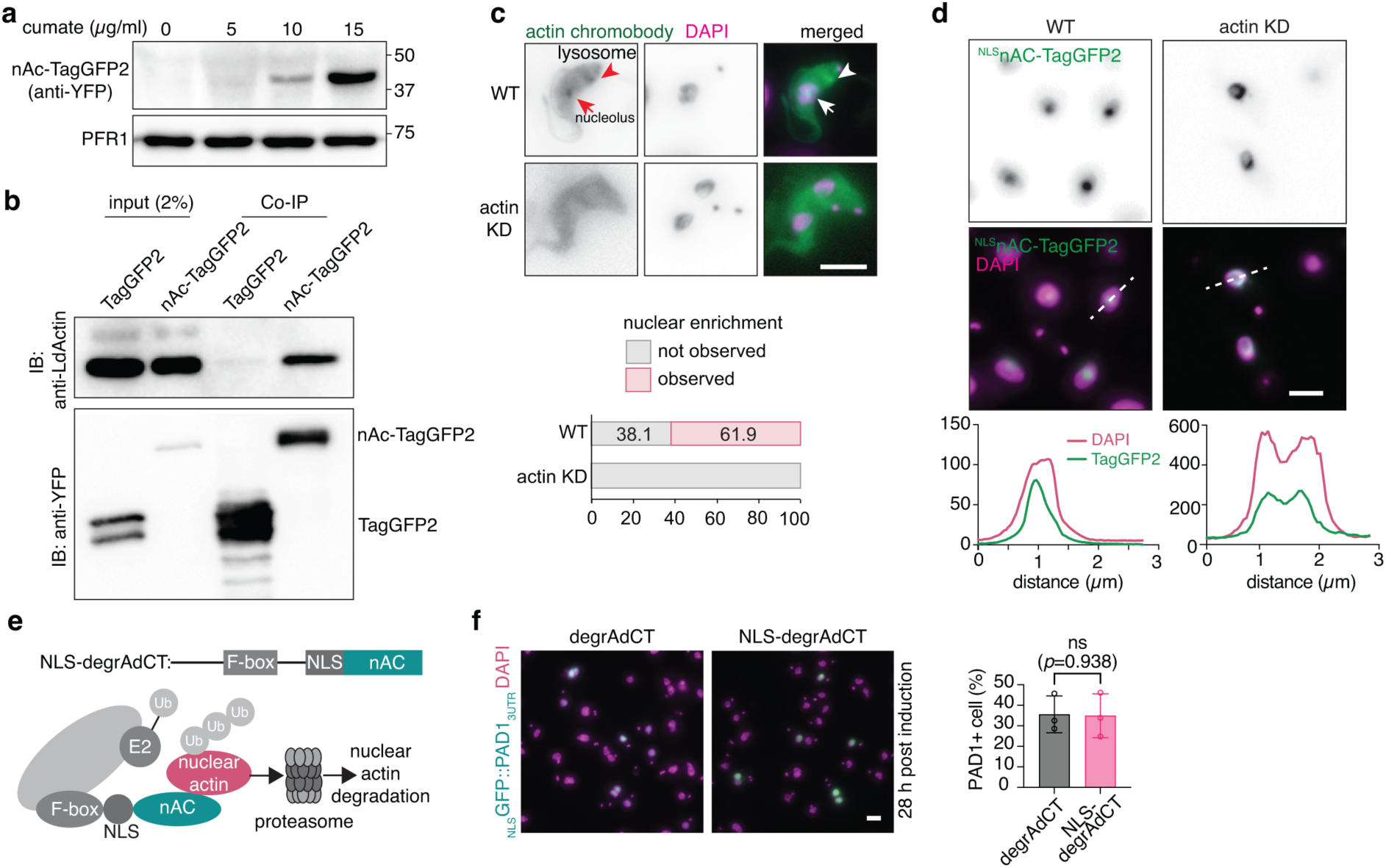
Degradation of nuclear actin triggers slender-to-stumpy differentiation. **a)** Cumate inducible expression of actin chromobody (nAc-TagGFP2) in BSF *T. brucei*. Cells were induced with 0, 5, 10 or 15 µg/ml cumate for 24 h and the expression of nAc-TagGFP2 was analyzed by immunoblotting with anti-GFP. PFR1 was used as loading control. **b)** nAc-TagGFP2 binds to *T. brucei* actin as shown by co-IP. Lysates of cells expressing nAc-TagGFP2 or TagGFP2 only were incubated with GFP-Trap beads. Co-precipitation of actin and nAc-TagGFP2 was detected using anti-actin and anti-GFP, respectively. **c)** nAc-TagGFP2 is present in the nucleus (arrow) and the lysosome (arrowhead) of BSF cells. Specificity of the labelling was confirmed using cells depleted of actin. Nuclear actin was observed in >60% of BSF cells. More than 200 cells were count in the quantification. Scale bar: 5 µm. **d)** The localization of nAc-TagGFP2 with a nuclear localization sequence (NLS) in BSF cells. The enrichment of ^NLS^nAc-TagGFP2 in the nucleolus in control cells is aborted upon actin depletion. The nucleolus distribution of actin was analyzed by plot profiles in control and actin KD cells. Scale bar: 5 µm. **e)** Diagram showing a modified degron system for targeted degradation of cytosolic or nuclear actin. using actin nAc (degrAdCT) or ^NLS^nAc (NLS-degrAdCT), respectively. **f)** PAD1-expression in cells induced for cytosolic or nuclear actin degradation. Quantitation of PAD1-positive cells at 28 h post actin degradation was shown as mean ± SD for 3 independent experiments. More than 200 cells were measured for each experiment. *p* values were calculated by 2 tailed, unpaired t-test using Prism GraphPad (ver. 10). Scale bar: 5 µm.

Nuclear actin has been shown to regulate RNA polymerase II (RNAP2)-mediated transcription through two different mechanisms^56,61^ (Fig. 7a). In one mechanism, actin functions as a component of the chromatin remodelling complex SWR1, which is involved in H2A.Z deposition and RNAP2 recruitment to the transcription starting site (TSS)^62^. Alternatively, actin can activate transcription process via direct interaction with RNA polymerase II^63,64^. As RNAP2 is responsible for the transcription of most protein-coding genes, a simple model would be that in the absence of actin, RNAP2 activity is inhibited, leading to cell quiescence and slender-to-stumpy differentiation.

**Fig. 7:**
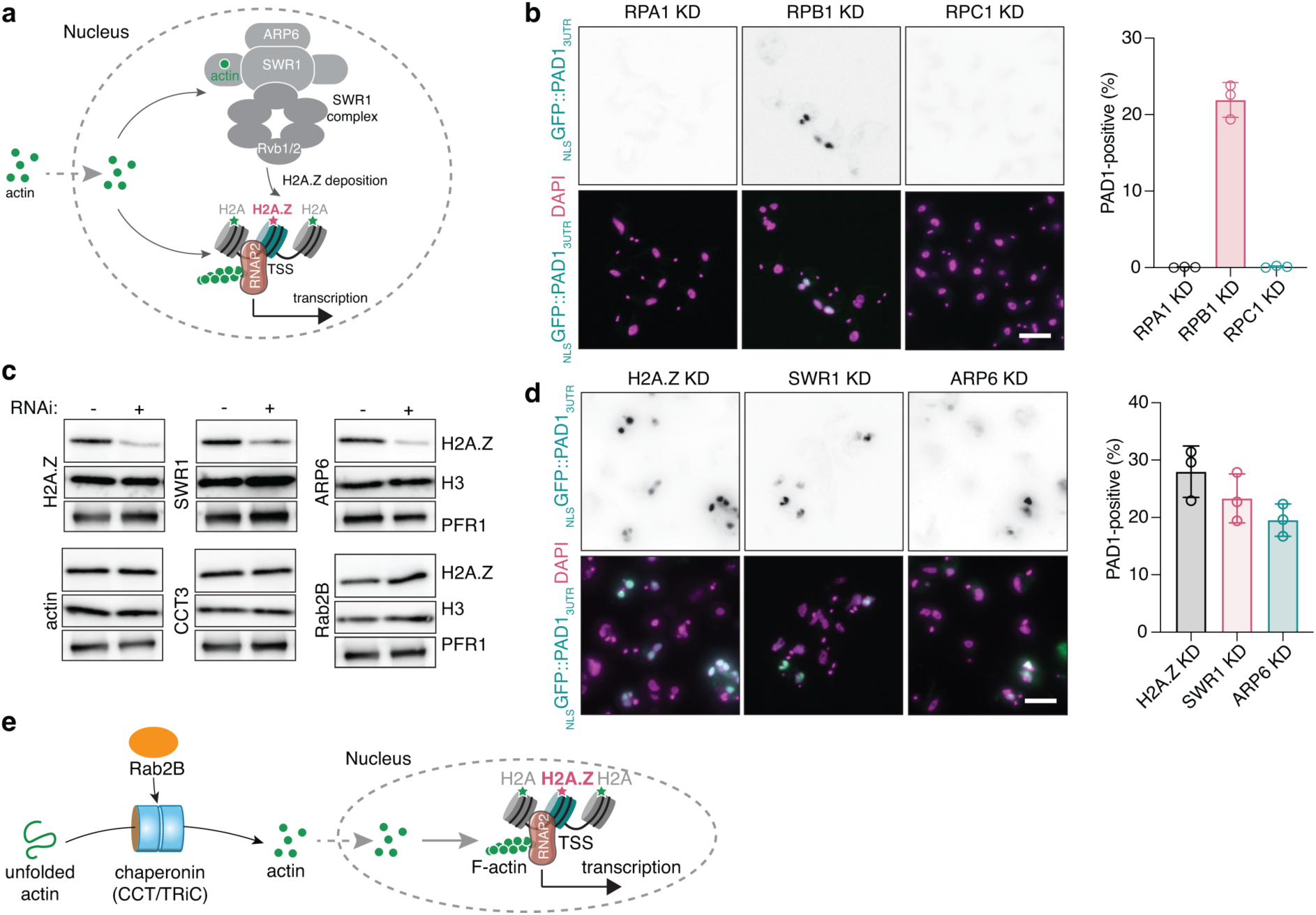
Actin KD-induced differentiation is independent of SWR1 complex-mediated chromatin remodelling. **a)** Schematic diagram showing two transcription regulation pathways that involve nuclear actin: 1) actin monomer as a component of SWR1 complex, which is required for H2A.Z deposition to the transcription starting site (TSS); and 2) actin filament that directedly regulates the inducible transcription by RNAP2. **b)** Knockdown of RNA polymerase II, but not I and III, triggers slender-to-stumpy differentiation. Differentiation was monitored by PAD1 expression in cells induced for RNAi targeting the largest subunit of RNA polymerase I (RPA1), II (RPB1) and III (RPC1). The quantification of PAD1-positive cells was displayed as mean ± SD of 3 independent experiments. More than 200 cells were measured for each experiment. Scale bar: 5 µm. **c)** RNAi of ARP6 and SWR1, subunits of the SWR1 complex, blocked H2A.Z deposition. RNAi of actin, CCT3, and Rab2B did not affect H2A.Z deposition. Specificity of anti-H2A.Z was confirmed by H2A.Z RNAi. PFR1 and H3 were used as loading control for the crude nuclear fractionation. **d)** RNAi depletion of SWR1 subunits ARP6 and SWR1, or H2A.Z triggered slender-to-stumpy differentiation. The quantification of PAD1-positive cells was displayed as mean ± SD of 3 independent experiments. More than 200 cells were quantitated in each experiment. Scale bar: 5 µm. **e)** Proposed working model for the function of Rab2B in slender-to-stumpy differentiation: Rab2B associates with chaperonin CCT/TRiC and regulates CCT/TRiC-mediated actin folding. Nuclear actin, likely in the form of actin filament, activates transcription mediated by RNAP2 and prevents cell differentiation.

To test the above possibilities, we first evaluated whether RNA polymerase activity contributes to the slender-to-stumpy differentiation. An RNAi screening against the largest subunits of each RNA polymerase complexes: RPA1 in RNAP1, RPB1 in RNAP2, and RPC1 in RNAP3 was performed, and we found that only RPB1 depletion induced slender-to-stumpy differentiation (Fig. 7b). To investigate the involvement of SWR1 complex and H2A.Z deposition in differentiation, we silenced the expression of two known components of the SWR1 complex, ARP6 and SWR1, as well as H2A.Z itself. Depletion of APR6 and SWR1 both reduced H2A.Z deposition at the TSS (Fig. 7c) and triggered cell differentiation (Fig. 7d). H2A.Z depletion also led to differentiation (Fig. 7d). To our surprise, although the depletion of Rab2B and actin both triggered differentiation, no effects on H2A.Z deposition were observed (Fig. 7c) in these cells. This is consistent with a recent report that actin was not found in the *T. brucei* SWR1 complex^65^. Together our results suggested a role of Rab2B and actin in transcriptomic remodelling through a pathway independent of chromatin remodelling or H2A.Z (Fig. 7e).

## DISCUSSION

In this study, we characterized the function of small GTPase Rab2B in autophagosome-lysosome fusion during *T. brucei* autophagy. We also discovered an unexpected role of Rab2B, CCT/TRiC, actin and RNA polymerase II in regulating slender-to-stumpy differentiation. Based on our results, we propose a mechanism of Rab2B in promoting chaperonin CCT/TRiC-mediated folding of actin, which further controls developmental transcription reprogramming and life cycle differentiation in *T. brucei*.

The highly conserved Rab2 GTPases are found in mammals, plants, and protozoa, but missing in a few organisms including budding yeast^66^. It has been proposed to function in ER-to-Golgi trafficking or in the formation of secretory granules based on its localization in the Golgi apparatus^67^. In a more recent study, Rab2 is shown to function in transversal T-tubule remodelling through genetic screening of a developmentally regulated remodelling program in *Drosophila* abdominal muscles^18^. This function is mediated by autophagy, during which Rab2 promotes autophagosome-lysosome fusion and further cargo degradation^18^. In mammalian cells, Rab2 recruits the homotypic fusion and protein sorting (HOPS) complex to autophagosome by interacting with HOPS subunit VPS39^26,68^. Assembly of functional HOPS complex between autophagosome and lysosome promotes membrane fusion driven by STX17-SNAP29-VAMP8^26^. In *T. brucei*, the HOPS subunits are well conserved but the SNARE proteins STX17 and SNAP29 are missing. As such, Rab2B function in autophagy appears conserved in T. brucei, the molecular mechanism however, maybe distinct to that found in other eukaryotes.

Interestingly, while Rab2B functioned in autophagy in both procyclic and bloodstream form trypanosomes, depletion of Rab2B had distinct effects. While depletion of Rab2B had no obvious growth phenotype in the insect stage (procyclic form), it was lethal in bloodstream form cells. Interesting still, Rab2B depletion triggered differentiation of proliferating slender cells to a quiescent stumpy form. The effect of Rab2B on cell differentiation is independent of autophagy, as inhibition of autophagy did not trigger differentiation nor inhibit Rab2B RNAi-induced differentiation. Consistently, in our searches for Rab2B effectors, we did not identify any candidates with known functions in autophagosome-lysosome fusion or autophagy.

The only Rab2B effector identified in this study is the chaperonin CCT/TRiC complex, which interacted with Rab2B in a GTP-dependent manner. In *Leishmania donovani*, a species closely related to *T. brucei*, a survey for proteins interacting with CCT/TRiC subunit γ identifies cytoskeletal proteins actin and α/β tubulin, several 7-bladed WD-domain containing proteins including Raptor and Gβ-like proteins as folding substrates^69^. Interestingly, three Rab small GTPases Rab1, Rab5 and Rab11, but not Rab2B, are also found in the CCT/TRiC-interactome^69^. Strikingly, depletion of CCT/TRiC subunits or well-known CCT/TRiC substrate actin, or treatment with actin polymerization inhibitor latrunculin A also triggered slender-to-stumpy differentiation, phenocopying Rab2B RNAi. These results suggested that Rab2B, CCT/TRiC and actin all function in the same pathway leading to cell differentiation. The simplest model to explain all of these observations is that Rab2B regulates CCT/TRiC-mediated actin folding and subsequent filament formation. The regulatory role of Rab GTPase to chaperonin function has also been reported during virus EV-A71 infection in mammalian cells, where Rab11 recruits chaperonin to assist the folding of individual virus structural proteins or inducing conformational change of newly formed EV-A71 provirons^70^. The regulatory role of Rab2B on actin filament is also reported in *Drosophila* where the F-actin (Myofibrils) organization is disrupted in the dorsal internal oblique muscles (IOMs) upon Rab2 RNAi^18^. In this study, we also demonstrated that Rab2B mutants affected actin filament organization in both *T. brucei* and mammalian cells. The precise mechanism of how Rab2B regulates CCT/TRiC-mediated actin folding remains to be investigated.

Although the functions of actin in cell motility, cell division and cytokinesis, cell signaling, vesicular trafficking, and the establishment and maintenance of cell junctions and cell shape are well established in various model organisms^71^, functional studies of actin in *T. brucei* and related parasites have been limited. A main challenge in actin studies in protozoan cells is the lack of microscopy tools to detect actin microfilaments although actin protein sequence is highly conserved. In *T. brucei,* no actin filament is detected by electron microscopy or staining with phalloidin^72^. Using an actin nanobody (nAC), we observed actin signal in the lysosome region as previously reported^59,73^, and additional enrichment in the nucleolus similar to those observed in mammalian cells and in *Leishmania*^74,75^.

The presence of nuclear actin is well established^76,77^ and an increasing body of evidence indicates that the nuclear actin is required to regulate RNAP2-mediated transcription, in the form of monomer^63^ or filament^55,78,79^. Actin has been shown to regulate transcription in at least two different ways. First, actin monomer functions as a component of chromatin remodelling SWR1 complex, which deposits the H2A variant H2A.Z to transcription starting sites (TSSs) for RNAP2 recruitment^80,81^.

Alternatively, F-actin directly interacts with and activate RNAP2^55,61,64^. In this study, we showed that depletion of RNAP2 or SWR1 subunits ARP6 (Tb927.10.2000, aka TbARP3) and SWR1 all led to cell differentiation phenotypes. However, depletion of Rab2B or actin did not affect H2A.Z deposition. The results suggested that actin-depletion induces transcriptional reprogramming and cell differentiation independent of chromosomal remodelling via the SWR1 complex, supporting recent observation that actin is not a component of trypanosome SWR1 complex as in mammals and yeast^82^. Given that LatA induced cell differentiation similar to actin depletion, *T. brucei* actin may regulate RNAP2 directly in filament form. While H2A.Z is a master regulator of cell differentiation and development through recruiting RNAP2, its regulation of RNAP2 transcription does not follow “one size fits all” but also dependents on various post-translational modifications (PTMs)^83^, diverse nucleosome composition and regulators such as actin.^84^ We thus propose that in *T. brucei,* actin-depletion causes transcription reprogramming and cell quiescence by regulating RNAP2 activity in SWR1 and H2A.Z-independent manner. As new players in slender-to-stumpy differentiation, Rab2B and actin shed new insights to the intricate transcriptional regulation of this process.

In addition to the SIF-induced differentiation pathway, alternative SIF-dependent pathway via adenosine monophosphate kinase (AMPK)/unconventional Target of Rapamycin (TbTOR4)^85,86^, and SIF-independent pathway via VSG transcriptional attenuation^45,46^ have also been reported. While each of these pathways could function during parasite differentiation in vivo and together ensure stumpy formation and parasite transmission, it is not clear if there is crosstalk among these pathways, and whether or how these different pathways all converge to the same cell fate determinants leading to stumpy formation. As ZC3H20, a key player in the SIF pathway, is also essential for Rab2B KD-induced differentiation, it is tempting to speculate that depletion of Rab2B or actin may bypass the SIF signalling mechanism that is missing in monophorphic strains, and induce differentiation through the SIF pathway. This study showed that Rab2B/actin-RNAi or LatA treatment readily triggered slender cell differentiation to stumpy and even procyclic forms, providing useful tools to study differentiation biology in lab strains.

## Materials and methods

### Reagents

LysoTracker Red DNA-99 (L7528, working concentration 10 µM), LysoSensor Green DND-189 (L7535, working concentration 10 µM) and MitoTracker Red CMXRos (M7512, working concentration 0.1 µg/ml) were purchased from Invitrogen (ThermoFisher Scientific). Latrunculin A (LatA, working concentration 0.2 µM) was purchased from Sigma (L5163). All other chemicals were purchased from Sigma.

### Cultivation of *Trypanosoma brucei* strains and HeLa cell

*T. brucei* procyclic (PCF) strains 29.13^87^, YTat1.1^88^ or DIY^43^ were cultivated in Cunningham medium^89^ supplemented with 10% Fetal Bovine Serum (FBS) at 27 °C. *T. brucei* bloodstream (BSF) strains Single Marker (SM)^87^, Lister 427^90^ or DI427^43^ were cultivated in HMI-9 medium^91^ supplemented with 10% FBS at 37 °C with 5% CO_2_. Mammalian HeLa cells were cultured in Dulbecco’s Modified Eagle Medium (DMEM), high glucose (Cat# 11965118, Thermo Scientific, US) supplemented with 10% FBS at 37 °C with 5% CO_2_.

### Vector construction and transfection

For inducible RNAi of Rab1A (Tb927.8.890), Rab1B (Tb927.9.15930), Rab2B (Tb927.11.15240), Rab4 (Tb927.11.13750), Rab5A (Tb927.10.12960), Rab5B (Tb927.11.4570), Rab6 (Tb927.2.2130), Rab7 (Tb927.9.11000), Rab7-like (Tb927.8.4620), Rab11 (Tb927.8.4330), Rab18 (Tb927.8.4770), Rab21 (Tb927.10.1520), Rab23 (Tb927.10.8560), Rab28 (Tb927.6.3040), RabX1 (Tb927.8.4610), RabX2 (Tb927.4.4220), ZC3H20 (Tb927.7.2660), HSP60 (Tb927.10.6400), PABP1 (Tb927.9.9290), PABP2 (Tb927.9.10770), CCT3 (Tb927.10.1060), CCT6 (Tb927.11.3240), GAT2 (Tb927.11.3130), ATG3 (Tb927.2.1890), Rab GDI (Tb927.3.4680), actin (A: Tb927.9.8850; B: Tb927.9.8880), α-tubulin (Tb927.1.2340/2360/2380/2400), β-tubulin (Tb927.1.2330/2350/2370/2390), Raptor (Tb927.11.620), mLST8 (Tb927.10.15640), CDC20 (Tb927.8.6500), H2A.Z (Tb927.7.6360), SWR1 (Tb927.11.10730), ARP6 (Tb927.10.2000), RPA1 (Tb927.8.5090), RPB1 (Tb927.4.5020/8.7400) and RPC1 (Tb927.10.2780), suitable fragments were selected using RNAit^92^, to avoid off-target effects. The fragments were amplified with the primers listed in Supplementary Table 1 and subcloned into p2T7 vectors^42^. For double RNAi, Rab2B fragment was cloned into BamHI/XhoI digested p2T7 vector. ZC3H20 or ATG3 fragments were further cloned into XbaI digested p2T7-Rab2B to generate the vectors for dKD.

For inducible expression of Rab2B, the full coding sequence of Rab2B was amplified, digested and cloned into pDEXCuo-mCherry vector^43^. The GDP- or GTP-bound S21N or Q66L mutants were created by point mutagenesis and confirmed by sequencing. For inducible expression of actin chromobody, the nAC-TagGFP2 fragment was amplified from vector Actin-Chromobody® Plasmid (TagGFP2) (ChromoTek, Germany) and cloned into HindIII/BamHI digested pDEXCuo-GFP/bla vector. To express the actin chromobody in nucleus, a nuclear leading sequence (NLS) from La protein^93^ was added to the N-terminus. To create the cytoplasmic or nuclear actin degron system, an XbaI restriction site was introduced into vector deGradFP and NLS-deGradFP^60^, respectively, and nAC (actin-VHH) was amplified and cloned into the vectors by replacing GFP nAb fragment to generate vectors degrAdCT and NLS-degrAdCT. Vector p4231 (courtesy of Mark Carrington), which consists of a *GFP* sequence with a nuclear localization signal that is followed by the 3’UTR of *PAD1*,^45^ was stably expressed in BSF cells and used as a stumpy marker.

To expression of GST-tagged Rab2B or actin nanobody (nAc) in bacteria, full sequence of Rab2B or nAc was amplified with primers listed in Supplementary Table 1 and cloned into BamHI/EcoRI or EcoRI/XhoI digested pGEX-6P-1 vector, respectively.

For transient expression of GFP-tagged HsRab2A/B in mammalian cells, the full sequence of HsRab2A/B was amplified from cDNA of 293T cells and cloned into vector pXJ40-GFP (courtesy of Liou Yih-Cherng). The GDP-bound (S20N) and GTP-bound (Q65L) mutants were created by point mutagenesis using primers listed in Supplementary Table 1.

### Autophagy assays

For amino acids starvation of PCF *T. brucei* cells, control or Rabs-RNAi cells harvested at mid-LOG phase were washed once with gHBSS (137 mM NaCl, 5.4 mM KCl, 0.25 mM Na_2_HPO_4_, 0.44 mM KH_2_PO_4_, 1.3 mM CaCl_2_, 1 mM MgSO_4_, 4.2 mM NaHCO_3_, 1 g/L (5.6 mM) glucose, pH 7.3) and resuspended in the same starvation buffer at a concentration of 5 × 10^6^ cells/ml and incubated at 27°C for 2 h unless otherwise indicated.

For amino acids starvation of BSF trypanosomes, control or Rab2B-RNAi cells harvested at mid-LOG phase were washed once with BSF cytomix (2 mM EGTA, pH 7.6, 120 mM KCl, 0.15 mM CaCl_2_, 10 mM K_2_HPO_4_/KH_2_PO_4_, pH 7.6, 25 mM HEPES, pH 7.6, 5 mM MgCl_2_, 0.5% glucose, 100 μg/ml BSA (Sigma, A7906), 1 mM hypoxanthine (Sigma, H9377), pH 7.6 and then resuspended in the same buffer at a concentration of 5 × 10^5^ cells/ml and incubated at 37 °C with 5% CO_2_ for 2 h.

After starvation, the cells were fixed with 4% PFA for 20 min at room temperature, washed with 1x PBS, and pelleted onto poly-L-lysine-coated coverslips by centrifugation. The formation of autophagosomes was monitored by fluorescence microscopy.

### *In vitro* differentiation of BSF trypanosomes to PCF

BSF cells were harvested by centrifugation at 1,800 *g* for 7 min at room temperature. Cells were washed with and resuspended in Cunningham medium at a concentration of 2-5 x 10^6^ cells/ml. After adding 6 mM tri-sodium citrate (Merck, 106448), cells were cultured at 27 °C. Cell differentiation was monitored by procyclin-expression with fluorescent microscopy or flow cytometry using anti-procyclin antibody (Cedarlane, Canada, CLP001A).

### Recombinant Rab2B overexpression, purification, on-beads digestion, and antibody preparation

The coding sequence of Rab2B (Tb927.11.15240) was amplified using primers indicated in Supplementary Table 1 and cloned into pGEX-6p-1 vector. The construct was then transformed into *E. coli* BL21 strain for expression. To increase the solubility of recombinant Rab2B, transformed BL21 cells were cultivated to OD600 ∼0.4. The cell culture was heat shocked at 42 °C for 10 min, followed by incubation at 37 °C for 20 min, 30 min on ice and another 20 min at 37 °C. Finally, the culture was induced with 0.1 mM IPTG for 20 hrs at 20 °C. The recombinant GST-tagged Rab2B was purified using Glutathione Sepharose 4B beads (GE Healthcare, USA) according to the manufacturer’s instruction. Rab2B protein was released from the beads by PreScission protease (GenScript, Z02779) treatment at 4 °C for 1 h. Purified Rab2B was dialyzed with PBS and used for customised antibody production in rats (SAbio, Singapore).

### Co-immunoprecipitation and mass spectrometry

The expression of YFP, YFP-Rab2B, YFP-Rab2B^S21N^ or YFP-Rab2B^Q66L^ in PCF cells was induced by 10 µg/ml cumate (4-Isopropylbenzoic acid, Sigma, 268402) for 24-48 h. A total of 2 x 10^9^ cells were harvested and washed with equal volume of cold PBS and resuspended in 1 ml lysis buffer (1 x PBS, supplemented with 1% NP40 and 2 x cOmplete, EDTA-free protease inhibitor cocktail (Roche, 11873580001)). Cell lysis was performed for 30 min on ice with several pipetting and vortex. The cell debris was pelleted by centrifugation at 21,000g for 10 min at 4 °C. The cleared supernatant was incubated with ChromoTek GFP-Trap® magnetic agarose beads (ChromoTek, Germany, gtma) for 2 h at 4 °C with rotation. After extensive washes, the captured proteins were eluted from the beads by boiling with SDS loading buffer and resolved by 6-20% gradient SDS-PAGE. After silver staining, the bands specifically co-immunoprecipitated with YFP-Rab2B^Q66L^ were sliced out and processed for mass spectrometry (Bioscience Research Centre, Nangyang Technological University, Singapore).

### Immunoblotting

Cell lysates (from ∼5 × 10^6^ cells) were fractionated by 12% or 6-20% SDS-PAGE. After blotting onto PVDF membranes (Bio-Rad, 162–0177), specific proteins were probed with rat-anti-TbRab2B (1:2,000, this study), rabbit-anti-LdCCT3 (1:2,000, courtesy of Neena Goyal), chicken-anti-LdActin (1:10,000, courtesy of Chhitar M. Gupta), rabbit-anti-trypanopain (1:2,000), rabbit-anti-H2A.Z and H3 (1:500 and 1:20,000, respectively, courtesy of Nicolai Siegel), rat-anti-TbATG3 (1:2,000), rabbit-anti-YFP (1:1,000, produced in our lab) or mouse-anti-p67 (1:2,000). Rabbit-anti-VDAC (1:2,000, courtesy of André Schneider)^94^ or rat-anti-PFR1 (1:2,500) were used as loading control for cytoplasmic or cytoskeletal proteins, respectively. Following incubation with secondary antibodies conjugated to horseradish peroxidase, the blots were visualized using the Clarity Western ECL system (Bio-Rad, 170– 5060) and documented using an ImageQuant LAS 4000 mini Luminescent Image Analyzer (GE Healthcare Life Sciences, Pittsburgh, PA, USA).

### Fluorescence microscopy

For procyclic form *T. brucei*, cells were attached onto cover slips and fixed with 4% PFA for 20 min. BSF cells were fixed with 4% PFA for 20 min then sedimented onto poly-L-lysine-coated cover slips by centrifugation. After permeabilization with 1% NP40 in PBS for 10 min and blocking with 3% BSA, the cells on the cover slips were stained with specified primary antibodies: anti-trypanopain (rabbit, 1:2000) and anti-p67 (mouse, 1:2,000) for the lysosomes, anti-GRASP (rabbit, 1:1,000) for the Golgi in *T. brucei*, anti-procyclin (mouse, 1:1,000), anti-GM130 (rabbit, 1:1,000) for the Golgi apparatus in mammalian cells. Nuclear and mitochondrial DNA was stained with 2 μg/ml DAPI (Invitrogen, D1306). Images were acquired using an Axioplan2 inverted microscope (Carl Zeiss MicroImaging, Jena, Germany) equipped with a CoolSNAP HQ2 camera (Photometrics, Tucson, Arizona, USA) and a Plan-Apochromat 63×/1.40 oil DIC objective.

To view actin distribution in mammalian HeLa cells, absolute ethanol-sterilized cover slips were place into 24-well plates and balanced with DMEM medium overnight. The HeLa cells were seeded onto the cover slips with a density of 3 x 10^5^ cells/ml. When cells achieved 60-70% confluence, co-transfections with mRFPRuby-LifeAct (Addgene plasmid #51009 originally from Rusty Lansford, a courtesy of Wu Min)^95^ and pXJ40-GFP-HsRab2A/B, pXJ40-GFP-HsRab2A/B^S20N^, or pXJ40-GFP-HsRab2A/B^Q65L^ were performed using Lipofectamine 3000 (Thermo Fisher Scientific, USA). At 24, 48 or 72 hours post transfection, the cells were fixed with 4% PFA, co-stained with DAPI and the expression and localization of LifeAct was monitored by fluorescence microscopy as described above.

### Transmission electron microscopy

Control or Rab2B RNAi cells were fixed by high pressure freezing (Technotrade International), freeze substituted with 1% OsO_4_ and 0.05% uranyl acetate in acetone in a Leica EM AF S2 system for 48 hours at −90 °C, 24 hours at −60 °C, 18 hours at −30 °C and 4 hours at 0 °C. Samples were then infiltrated with LX112 resin (Ladd Research Industries, USA), polymerized, cut into 120 – 200 nm sections and examined with an Tecnai T12 transmission electron microscope (FEI, USA) operating at 120 kV.

### RNA-Seq

The total RNAs were isolated from 5×10^7^ cells with RNeazy® mini Kit (QIAGEN, Germany) and treated with RNase free DNase I (EN0521, Thermo Scientific, USA). The library was prepared using Illumina Stranded mRNA Library Preparation Kit. Quality and concentration of all libraries was determined by capillary electrophoresis and sequenced (one lane, paired end 150 bp) with Illumina Hiseq 2000 System (Illumina, USA). Each sample gives around 25 million paired end reads (unfiltered). Transcriptome sequences were extracted from the 11 megabase chromosomes of *T. brucei* TREU927 reference genome using gffread version 0.11.8 with the corresponding genome annotation file, and read counts for the annotated genes were quantified against this reference transcriptome using Salmon version 1.2.1 with quasi-mapping-based mode and default parameters. Transcript abundances were reported as transcript per million (TPM) values. Differential gene expression was conducted using iDEP (version 0.91) online analyses (http://bioinformatics.sdstate.edu/idep/)^96^ with DEseq2 method and default parameters by comparing Rab2B-depleted cells (in triplicate) to Lister 427 cells (in duplicate), actin-depleted cells (in triplicate) to Lister 427cells, Rab2B-depleted cells to actin-depleted cells or newly differentiated PCF cells (in duplicate) to Lister 427 cells. The cut-off point was set to 0.1 for FDR and the minimum fold change shown was 2.

### Lysosome function assay

To track the LysoTracker Red DND-99 (Molecular Probes, L7528) uptake, control or Rab2B-KD BSF cells were cultivated in HMI-9 medium to a density of 1 x 10^6^ cells/ml and stained with 10 µM LysoTracker Red for 30 min at 37 °C (the Rab2B-KD live cells were co-stained with 5 µg/ml DAPI). After washing twice with medium or PBSG (PBS supplemented with 1 g/L glucose), cells were fixed with 4% PFA for 20 min at room temperature. After washing twice with 1x PBS, the control and Rab2B-KD cells were mixed 1:1 and sedimented onto poly-L-lysine-coated coverslips by centrifugation.

To tracking the LysoSensor staining in live cell, control and Rab2B-KD cells were stained with 10 µM LysoSensor Green DND-189 (Molecular Probes, L7535) in medium for 30 min (the Rab2B-KD cells were co-stained with 5 µg/ml DAPI). After washing with PBSG for three times, control and Rab2B-KD cells were mixed 1:1, smeared on glass slides and imaged with fluorescence microscopy immediately.

To track the maturation of cathepsin L, the cell lysates of control or Rab2B-KD cells were resolved by 6-20% SDS-PAGE and the cathepsin protein was probed with anti-trypanopain antibody.^97^ The three bands (from top to bottom) represent immature, intermediate and mature form of cathepsin, respectively.

### Mitochondria staining

Control or Rab2B-KD BSF cells were stained with 0.1 μg/ml MitoTracker Red CMXRos (Molecule Probes, M7512) for 30 min at 37 °C. Cells were washed twice with PBSG, fixed with 4% PFA, permeablized with 1% NP-40, stained with DAPI, sedimented onto poly-L-lysine-coated cover slips by centrifugation and imaged by fluorescence microscopy.

### Extraction of chromatin-associated proteins

The chromatin extraction protocol described previously was used to analyse the amount of chromatin-associated histones^83^. Briefly, cells were harvested by centrifugation (1,500 × *g* for 10 min at 4 °C) and washed in 1 ml of trypanosome dilution buffer (TDB: 5 mM KCl, 80 mM NaCl, 1 mM MgSO_4_, 20 mM Na_2_HPO_4_, 2 mM NaH_2_PO_4_, 20 mM glucose, pH 7.4). The cell pellet was solubilized in CSK-buffer (100 mM NaCl, 0.1% Triton X-100, 300 mM sucrose, 1 mM MgCl_2_, 1 mM EGTA, 10 mM PIPES (pH 6.8; with NaOH) supplemented with *cOmplete* Protease Inhibitor Cocktail (Roche) for 10 min at 4 °C. The insoluble fraction was separated with centrifugation at 2,550 *g* for 5 min at 4 °C. The pellet was washed with CSK-buffer, resuspended in SDS-loading buffer and boiled at 90 °C for 10 min. The proteins were separated via SDS-PAGE and analysed by immunoblotting.

### RNA isolation and cDNA synthesis from mammalian 293T cells

The total RNA from about 3 x 10^6^ 293T cells were isolated using TRIzol reagent (Life Technologies, 15596-026) following the protocol provided. After treatment with RNase-free DNase I (Roche, 04 716 728 001), the first-strand cDNA was synthesized with M-MLV Reverse Transcriptase (Invitrogen, 28025-013) following the provided protocol.

### Bioinformatic analyses

The protein sequences of all eight CCT/TRiC subunits was atrieved from TriTrypDB (https://tritrypdb.org/tritrypdb/app) based on the annotation and was further confirmed by NCBI protein BLAST. The orthologues from other model organisms were atrieved by BLAST against Model Organisms (landmarks) database using query sequences of human CCT/TRiC subunits. Multiple protein sequence alignment and phylogenetic analysis was conducted using Clustal Omega from EMBL-EBI (https://www.ebi.ac.uk/jdispatcher/msa/clustalo). The phylogenetic tree was displayed and edited using iTOL (https://itol.embl.de).

### Statistical analyses

For all quantifications, 3 independent experiments were performed and statistical analyses were performed using the unpaired and 2-tailed, equal variance Student *t* test using GraphPad Prism 10 for Mac OS (GraphPad Software, Boston, Massachusetts USA, www.graphpad.com). *P* values <0.05 were determined to be statistically significant, while > 0.05 was determined as not statistically significant (NS). Exact *P* value numbers were shown on the graph if greater than 0.0001, otherwise shown as *P*<0.0001.

## Data Availability

RNA-Seq raw files have been deposited to the NCBI Sequence Read Archive (SRA) under BioProject accession number PRJNA1184179 (sample numbers SAMN44665644-SAMN44665653). The processed files have been deposited to the NCBI Gene Expression Omnibus (GEO) with accession number GSE282119. Source data are provided with this paper.

## ACKNOWLEDGEMENTS

We thank Mark Carrington from University of Cambridge and Wu Min from Yale School of Medicine for providing vectors p4231 and mRFPRuby-LifeAct, respectively. We also thank Chhitar M Gupta from Institute of Bioinformatics and Applied Biotechnology, India, Neena Goyal from CSIR-Central Drug Research Institute, India, Nicolai Siegel from Ludwig-Maximilians-Universität, Germany, and André Schneider from Department of Chemistry and Biochemistry, University of Bern, Switzerland for providing us the antibodies against LdActin, LdCCTγ, H2A.Z/H3 and TbVDAC, respectively. We also thank Tan Lay Keng Angie from Duke-NUS, Singapore for RNA-Seq, Meng Wei from Nangyang Technological University, Singapore for mass spectrometry analyses and Wang Xiaoning from School of Medicine, National University of Singapore for flow cytometry analyses. This research was funded by Ministry of Education, Singapore (MOE-T2EP30221-0002 and A-0008403-00-00) to CYH.

## AUTHOR CONTRIBUTIONS

F-J.L. and C.Y.H. conceived and designed the project; F-J.L. conducted most of the experiments and analyses; X.D. contributes to electronic microscopy analysis; Y.H. contributes to co-immunoprecipitation experiment; D.H. & E.C. contribute to RNA-Seq analysis; F-J.L. and C.Y.H. wrote the manuscript.

---

**Supplementary Fig. 1:**

**a)** Growth analyses of PCF cells induced for Rab1A RNAi or not. Only 1 experiment was performed and displayed.

**b)** Rab1A depletion inhibited starvation-induced autophagy in PCF cells. Number of autophagosomes per cell was shown as mean ± SD of 3 independent experiments. More than 200 cells were measured for each experiment. Scale bar: 5 µm.

**c)** Growth analyses of PCF cells induced for Rab11 RNAi or not. Only 1 experiment was performed and displayed.

**d)** Increased autophagosome in Rab11 KD cells, under fed or starvation conditions. Number of autophagosomes per cell was shown as mean ± SD of 3 independent experiments. More than 200 cells were measured for each experiment. Scale bar: 5 µm.

**e)** Rab11 depletion inhibited cytokinesis and generated multinucleated cells. Although the number of autophagosomes/cell increased upon Rab11 RNAi, autophagosome number/nucleus did not change, suggesting that the increase in autophagosome number was linked to cytokinesis arrest.

**f)** In PCF cells, Rab2B KD had no obvious effect on Golgi (marked with anti-GRASP), but caused the increasing of lysosome number (marked with anti-p67). Scale bar: 5 µm.

**g)** In PCF cells, overexpression of Rab2BS^21N^ mutant phenocopied Rab2B KD, causing lysosome number increasing. Scale bar: 5 µm.

**h)** In BSF cells, Rab2B KD also led to multiple lysosomes as observed with lysosome membrane marker p67 and matrix marker trypanopain. Scale bar: 5 µm.

**i)** Schematic diagram showing the maturation and transport of p67 in BSF cells. p67 is synthesized as a 100 kDa ER species (gp100), which was converted to gp150 in the Golgi apparatus via post-translational modifications. The gp150 form was secreted to cell surface and further internalized into the lysosome and processed to gp75, gp42 and gp32 forms.

**j)** Rab2B KD affected the maturation of p67, leading to the accumulation of gp150 and decrease in gp75 and gp42 mature forms.

**k)** The accumulation of p67 on the cell surface in Rab2B KD cells. The cells were fixed with 4% PFA, with or without permeabilization by 1% NP-40 and stained with anti-p67 antibody. Distribution of the p67 signal was shown by plot profiles. Scale bar: 5 µm.

**Supplementary Fig.2:**

**a)** BSF cells were induced for Rab2B RNAi to trigger slender-to-stumpy differentiation. The stumpy cells were further differentiated to PCF cells. RNA-seq was performed and the total number of up- and down-regulated genes (cut-off fold change: ± 2) in newly differentiated PCF cells compared with parental BSF cells was shown as a bar graph.

**b)** PCA analyses showing transcription variation between the newly differentiated PCF and parental BSF cells.

**c)** A volcano plot showing differentially expressed transcripts in newly differentiated PCF and parental BSF cells. The top hits were highlighted.

**d)** Representative GO terms analyses of the transcriptomes of newly differentiated PCF cells compared with parental BSF cells. Genes involved in TCA cycle/cell respiration were upregulated and those involved in glucose metabolism/glycolysis were downregulated, consistent with the energy metabolism characteristics of PCF and BSF cells.

**Supplementary Fig. 3: Rab2B KD-triggered differentiation is autophagy independent.**

**a)** Immunoblotting shown the RNAi efficiency of Rab2B and ATG3 double knockdown after 24 h of induction, in BSF trypanosomes. Actin and PFR1 was used as loading control.

**b)** Fluorescent microscopy checking the expression of PAD1 in WT, Rab2B KD, ATG3 KD or double KD cells. The quantification was shown as mean ± SD for 3 independent experiments. More than 200 cells were measured for each experiment of each sample. *p* values were calculated by 2 tailed, unpaired t-test using Prism GraphPad (ver. 10). Scale bar: 5 µm.

**Supplementary Fig. 4:**

**a)** Phylogenetic analysis of CCT/TRiC subunits from *T. brucei*, *Homo sapiens*, *Mus Musculus*, *Danio rerio*, *Drosophila melanogaster*, *Caenorhabditis elegans*, *Arabidopsis thaliana*, *Dictyostelium discoideum, Saccharomyces cerevisiae, Schizosaccharomyces pombe, Plasmodium falciparum and Leishmania donovani.* Multiple protein sequence alignment and phylogenetic analysis was conducted using Clustal Omega from EMBL-EBI. The phylogenetic tree was displayed and edited using iTOL.

**b)** The identity (%) of amino acids sequence among CCT/TRiC subunits from *T. brucei* and human (*Homo sapiens*).

**c)** Predicted structure of *T. brucei* CCT1-8 subunits using trRosetta. The TM score reflected the similarity of *T. brucei* protein to its ortholog in *Homo sapiens*.

**Supplementary Fig. 5:** Representative GO term analyses of upregulated genes in BSF cells induced for Rab2B KD and actin KD, compared to control BSF cells.

**Supplementary Fig. 6: CCT/TRiC-mediated actin folding did not contribute to functional autophagy**

CCT6 knockdown or LatA treatment did not affect the starvation induced autophagy in PCF cells. Autophagosomes were labelled with YFP-ATG8.2 and number of phagosomes/cell was shown as 3 independent experiments. More than 200 cells were measured for each experiment. *p* values were calculated by 2 tailed, unpaired t-test using Prism GraphPad (ver. 10). Scale bar: 5 µm.

**Supplementary Fig. 7:**

**a)** Overexpression of S20N or Q66L mutants of HsRab2A or HsRab2B affected Golgi organization in HeLa cell. Scale bar: 10 µm.

**b)** Overexpression of HsRab2A variants did not affect actin organization in HeLa cells. Actin filaments were labelled with mRFPRuby-tagged LifeAct. Scale bar: 10 µm.

