## Supplementary Fig. 1-7; Supplementary Table 1 for "Rab2B promotes chaperonin-mediated actin folding and prevents developmental transcription reprogramming and quiescence in trypanosomes"

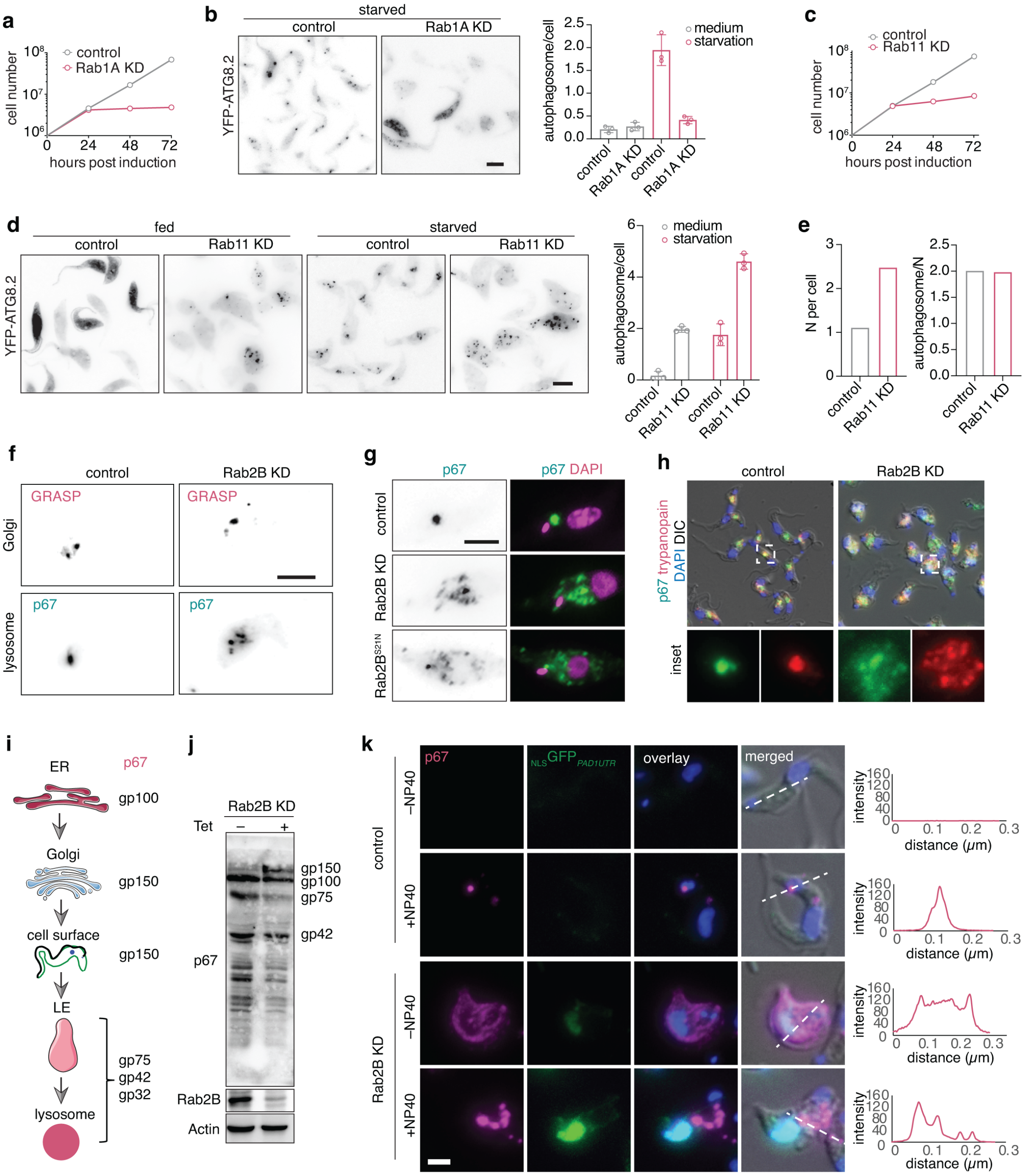


**Supplementary Fig. 1: a** Growth analyses of PCF cells induced for Rab1A RNAi or not. Only 1 experiment was performed and displayed. **b** Rab1A depletion inhibited starvation-induced autophagy in PCF cells. Number of autophagosomes per cell was shown as mean ± SD of 3 independent experiments. More than 200 cells were measured for each experiment. Scale bar: 5 μm. **c** Growth analyses of PCF cells induced for Rab11 RNAi or not. Only 1 experiment was performed and displayed. **d** Increased autophagosome in Rab11 KD cells, under fed or starvation conditions. Number of autophagosomes per cell was shown as mean ± SD of 3 independent experiments. More than 200 cells were measured for each experiment. Scale bar: 5 μm. **e** Rab11 depletion inhibited cytokinesis and generated multinucleated cells. Although the number of autophagosomes/cell increased upon Rab11 RNAi, autophagosome number/nucleus did not change, suggesting that the increase in autophagosome number was linked to cytokinesis arrest. **f** In PCF cells, Rab2B KD had no obvious effect on Golgi (marked with anti-GRASP), but caused the increasing of lysosome number (marked with anti-p67).

Scale bar: 5 μm. **g** In PCF cells, overexpression of Rab2BS$^{21N}$ mutant phenocopied Rab2B KD, causing lysosome number increasing. Scale bar: 5 μm. **h** In BSF cells, Rab2B KD also led to multiple lysosomes as observed with lysosome membrane marker p67 and matrix marker trypanopain. Scale bar: 5 μm. **i** Schematic diagram showing the maturation and transport of p67 in BSF cells. p67 is synthesized as a 100 kDa ER species (gp100), which was converted to gp150 in the Golgi apparatus via post-translational modifications. The gp150 form was secreted to cell surface and further internalized into the lysosome and processed to gp75, gp42 and gp32 forms. **j** Rab2B KD affected the maturation of p67, leading to the accumulation of gp150 and decrease in gp75 and gp42 mature forms. **k** The accumulation of p67 on the cell surface in Rab2B KD cells. The cells were fixed with 4% PFA, with or without permeabilization by 1% NP-40 and stained with anti-p67 antibody. Distribution of the p67 signal was shown by plot profiles. Scale bar: 5 μm.

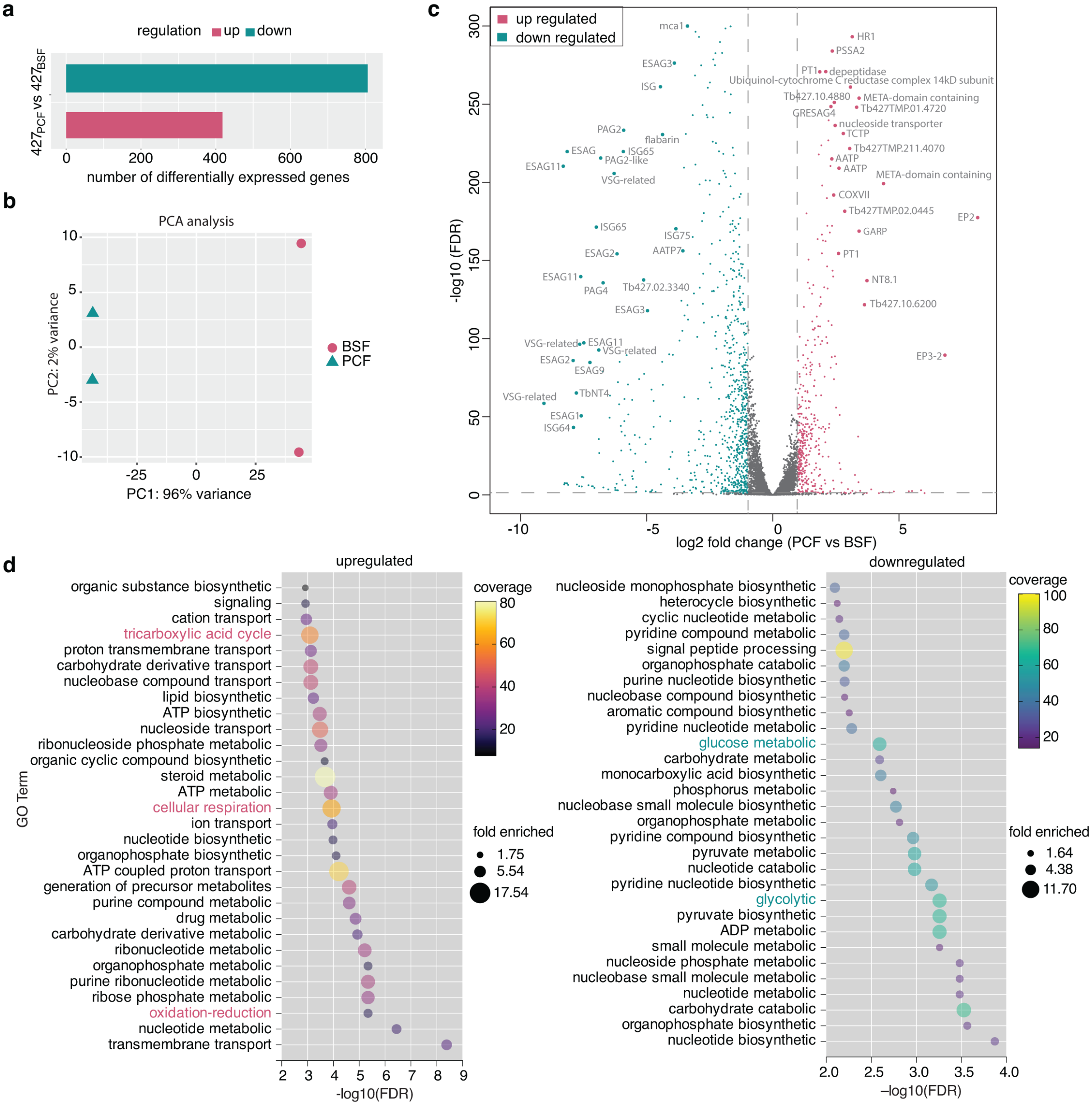


**Supplementary Fig. 2: a** BSF cells were induced for Rab2B RNAi to trigger slender-to-stumpy differentiation. The stumpy cells were further differentiated to PCF cells. RNA-seq was performed and the total number of up- and down-regulated genes (cut-off fold change: ± 2) in newly differentiated PCF cells compared with parental BSF cells was shown as a bar graph. **b** PCA analyses showing transcription variation between the newly differentiated PCF and parental BSF cells. **c** A volcano plot showing differentially expressed transcripts in newly differentiated PCF and parental BSF cells. The top hits were highlighted. **d** Representative GO terms analyses of the transcriptomes of newly differentiated PCF cells compared with parental BSF cells. Genes involved in TCA cycle/cell respiration were upregulated and those involved in glucose metabolism/glycolysis were downregulated, consistent with the energy metabolism characteristics of PCF and BSF cells.

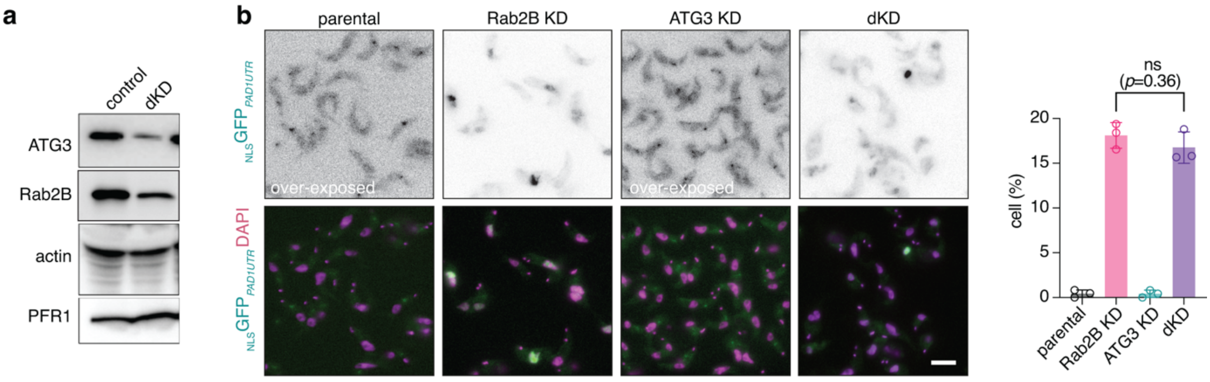


**Supplementary Fig. 3: Rab2B KD-triggered differentiation is autophagy independent. a** Immunoblotting shown the RNAi efficiency of Rab2B and ATG3 double knockdown after 24 h of induction, in BSF trypanosomes. Actin and PFR1 was used as loading control. **b** Fluorescent microscopy checking the expression of PAD1 in WT, Rab2B KD, ATG3 KD or double KD cells. The quantification was shown as mean ± SD for 3 independent experiments. More than 200 cells were measured for each experiment of each sample. *p* values were calculated by 2 tailed, unpaired t-test using Prism GraphPad (ver. 10). Scale bar: 5 µm.

**a**

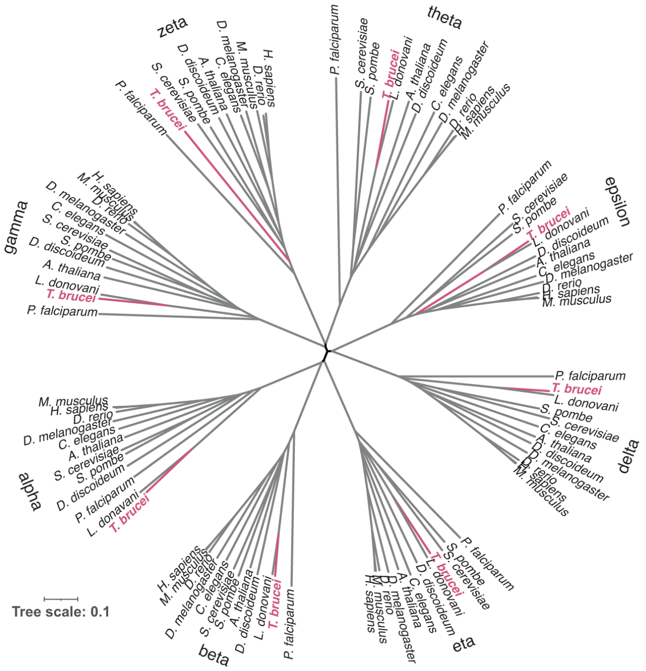


**b**

| identity (%) | *T. bruce*i CCT/TRiC subunits 1 | 2 | 3 | 4 | 5 | 6 | 7 | 8 |
|---|---|---|---|---|---|---|---|---|
| *H. sapiens* CCT/TRiC subunits 1 | 56.54 | 35.20 | 30.71 | 33.53 | 30.31 | 27.29 | 36.07 | 29.94 |
| 2 | 32.63 | 61.81 | 28.9 | 29.41 | 30.98 | 25.00 | 33.65 | 26.92 |
| 3 | 31.12 | 29.62 | 52.76 | 30.54 | 30.42 | 26.57 | 29.76 | 28.76 |
| 4 | 33.86 | 32.55 | 32.43 | 57.47 | 34.48 | 28.43 | 32.68 | 29.61 |
| 5 | 32.44 | 34.36 | 33.08 | 35.52 | 60.15 | 25.19 | 31.02 | 27.50 |
| 6 | 29.73 | 27.68 | 29.07 | 26.98 | 28.82 | 54.80 | 27.63 | 22.94 |
| 7 | 33.33 | 33.14 | 33.15 | 30.84 | 31.21 | 27.67 | 61.30 | 25.95 |
| 8 | 27.67 | 28.08 | 24.72 | 29.35 | 29.22 | 22.69 | 27.92 | 50.19 |

**c**

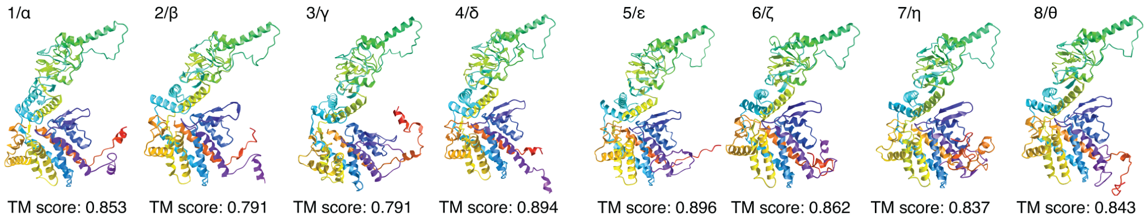


**Supplementary Fig. 4: a** Phylogenetic analysis of CCT/TRiC subunits from *T. brucei*, *Homo sapiens*, *Mus Musculus*, *Danio rerio*, *Drosophila melanogaster*, *Caenorhabditis elegans*, *Arabidopsis thaliana*, *Dictyostelium discoideum, Saccharomyces cerevisiae, Schizosaccharomyces pombe, Plasmodium falciparum and Leishmania donovani.* Multiple protein sequence alignment and phylogenetic analysis was conducted using Clustal Omega from EMBL-EBI. The phylogenetic tree was displayed and edited using iTOL. **b** The identity (%) of amino acids sequence among CCT/TRiC subunits from *T. brucei* and human (*Homo sapiens*). **c** Predicted structure of *T. brucei* CCT1-8 subunits using trRosetta. The TM score reflected the similarity of *T. brucei* protein to its ortholog in *Homo sapiens*.

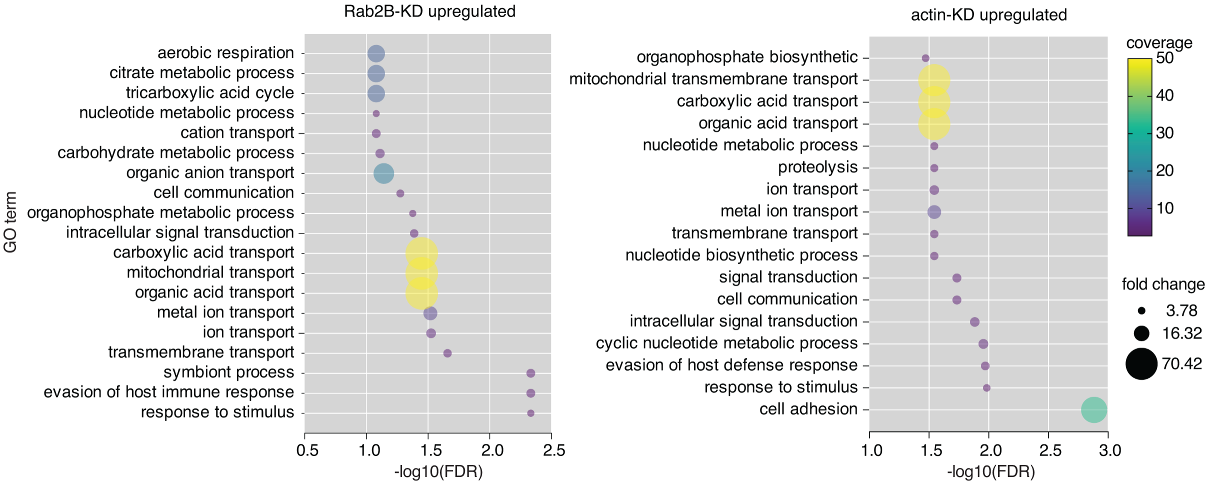


**Supplementary Fig. 5:** Representative GO term analyses of upregulated genes in BSF cells induced for Rab2B KD and actin KD, compared to control BSF cells.

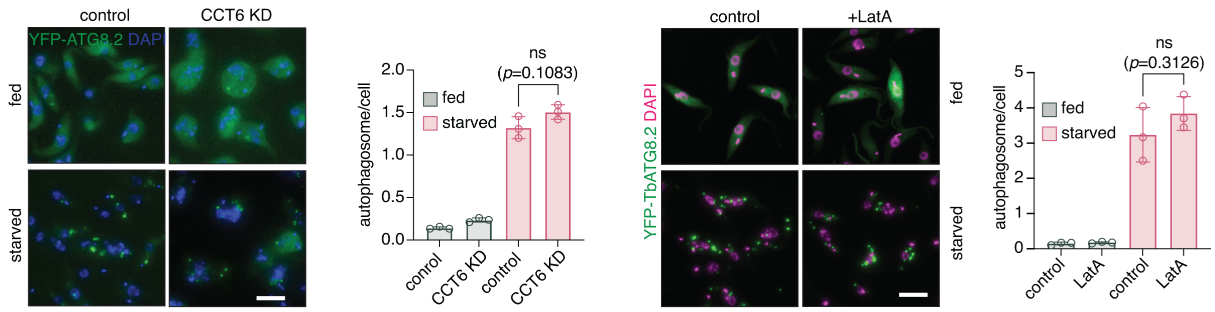


**Supplementary Fig. 6: CCT/TRiC-mediated actin folding did not contribute to functional autophagy.** CCT6 knockdown or LatA treatment did not affect the starvation induced autophagy in PCF cells. Autophagosomes were labelled with YFP-ATG8.2 and number of phagosomes/cell was shown as 3 independent experiments. More than 200 cells were measured for each experiment. *p* values were calculated by 2 tailed, unpaired t-test using Prism GraphPad (ver. 10). Scale bar: 5 μm.

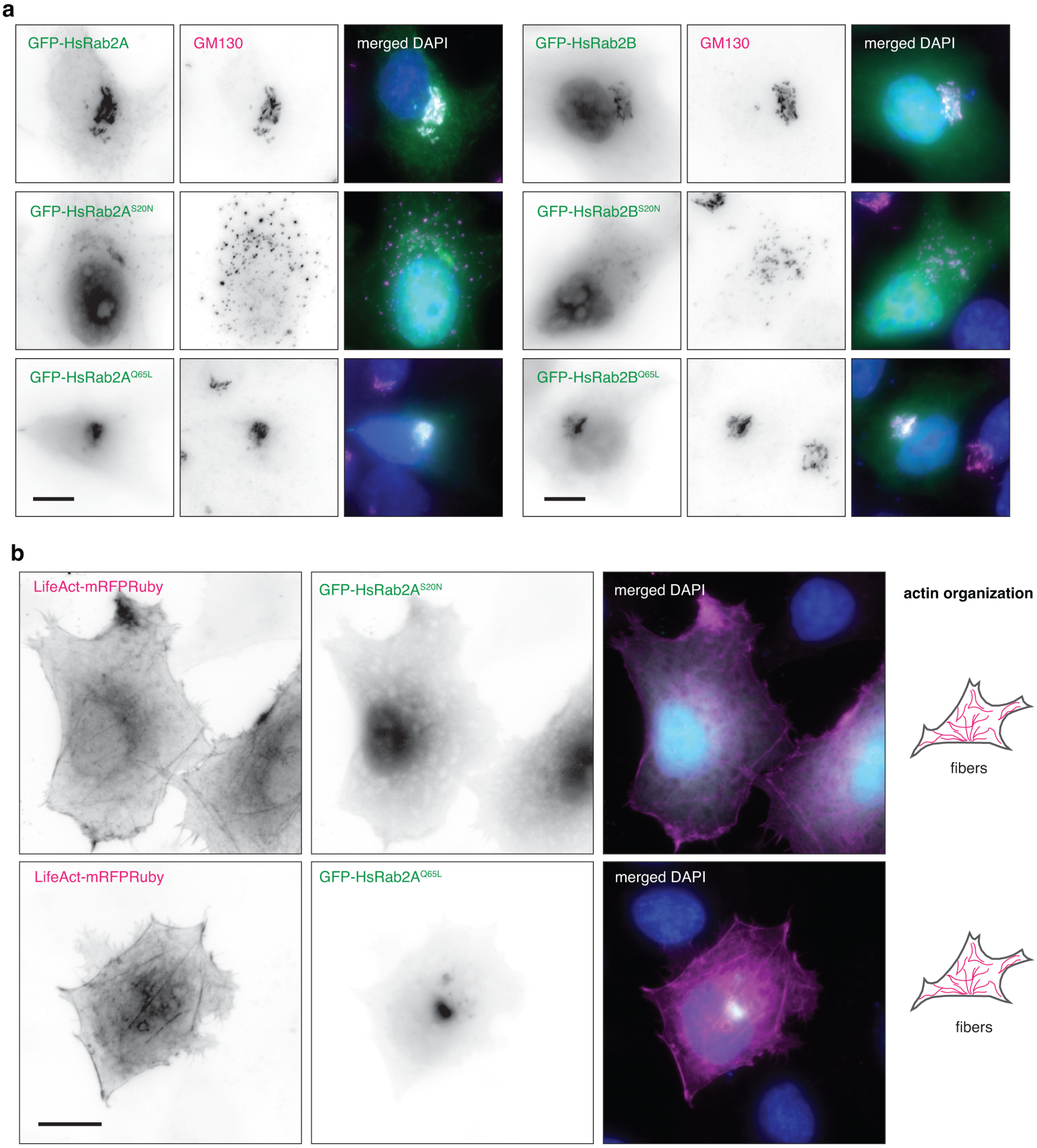


**Supplementary Fig. 7: a** Overexpression of S20N or Q66L mutants of HsRab2A or HsRab2B affected Golgi organization in HeLa cell. Scale bar: 10 μm. **b** Overexpression of HsRab2A variants did not affect actin organization in HeLa cells. Actin filaments were labelled with mRFPRuby-tagged LifeAct. Scale bar: 10 μm.

**Supplementary Table 1: a list of primers used in this study.**

| Primer name | Sequence (5' – 3') | Restriction site | purpose |
|---|---|---|---|
| Rab1A-iF | cgggatccgacgacagctacacggagag | BamHI | RNAi of Rab1A |
| Rab1A-iR | ccgctcgagagtaatggccttccaccagc | XhoI | |
| Rab1B-iF | gctctagaagcacagttctaccgtgacg | XbaI | RNAi of Rab1B |
| Rab1B-iR | gctctagatctctggctcctctcagacc | XbaI | |
| Rab2B-iF | cgggatccagcggagtgggaaagagttg | BamHI | RNAi of Rab2B |
| Rab2B-iR | ccgctcgagtcagaaacgcgtcatccaca | XhoI | |
| Rab4-iF | gctctagatgttggagatagcggcacag | XbaI | RNAi of Rab4 |
| Rab4-iR | gctctagaggtggaagcagacgtctcaa | XbaI | |
| Rab5A-iF | gctctagaacaggatgcaataaccgcca | XbaI | RNAi of Rab5A |
| Rab5A-iR | gctctagaagagccgggacctgaatttg | XbaI | |
| Rab5B-iF | gctctagacggtgttggtaaatcatccctcg | XbaI | RNAi of Rab5B |
| Rab5B-iR | gctctagaaacacaccaccgctatgaacag | XbaI | |
| Rab6-iF | gctctagacgccagttgcatccacaaaa | XbaI | RNAi of Rab6 |
| Rab6-iR | gctctagatttccactacgggaatggcc | XbaI | |
| Rab7-iF | gctctagacaaccggtacaaggcgacg | XbaI | RNAi of Rab7 |
| Rab7-iR | ccgctcgagcacaccaggcttctgccg | XhoI | |
| Rab7L-iF | gctctagagcaacatctcatcgcggttc | XbaI | RNAi of Rab7-like |
| Rab7L-iR | gctctagacgaagtagcacaagcaaccg | XbaI | |
| Rab11-iF | cgggatcctcatgactcgctacacagccg | BamHI | RNAi of Rab11 |
| Rab11-iR | ccgctcgagacgtttgttttctccaacgcg | XhoI | |
| Rab18-iF | gctctagattctctcgttctcgttgggc | XbaI | RNAi of Rab18 |
| Rab18-iR | gctctagaacctcatcaaaggcctgctc | XbaI | |
| Rab21-iF | gctctagaacagcagtgtcaacgttcca | XbaI | RNAi of Rab21 |
| Rab21-iR | gctctagaggctccactggcactattgt | XbaI | |
| Rab23-iF | gctctagatgatgggaatgttgggaaatcgtc | XbaI | RNAi of Rab23 |

| | | | |
|---|---|---|---|
| Rab23-iR | gctctagaccttcgtagagacgcgaaacaaag | XbaI | |
| Rab28-iF | gctctagatggcgctgttggtaagacat | XbaI | RNAi of Rab28 |
| Rab28-iR | gctctagaaaattagcaacgacgcgctc | XbaI | |
| RabX1-iF | gctctagagagcggtttcagtcagtgg | XbaI | RNAi of RabX1 |
| RabX1-iR | gctctagaccttcgtttccggctaacct | XbaI | |
| RabX2-iF | gctctagagcacttaacatgggggacga | XbaI | RNAi of RabX2 |
| RabX2-iR | gctctagactgtcacccttcgaaccgaa | XbaI | |
| ZC3H20-iF | cccaagcttgatattacgctctgccgcct | HindIII | RNAi of ZC3H20 or double RNAi of Rab2B and ZC3H20 |
| ZC3H20-iR | cccaagctttgccgtgaaagcgaaaaagg | HindIII | |
| HSP60-iF | cccaagctttgtaaccgttgcgaagtcga | HindIII | RNAi of HSP60 |
| HSP60-iR | cccaagcttcgaacgcatcttcaagctcg | HindIII | |
| PABP1-iF | cccaagcttcttttcgaagtttggcggca | HindIII | RNAi of PABP1 |
| PABP1-iR | cccaagctttgacaacgagaggtttcccg | HindIII | |
| PABP2-iF | cccaagcttctcgccctctgtttgtgagt | HindIII | RNAi of PABP2 |
| PABP2-iR | cccaagctttctggagtcatgctggcaag | HindIII | |
| CCT3-iF | gctctagacaaagtgaccggccaaaagg | XbaI | RNAi of CCT3 |
| CCT3-iR | gctctagagtatctttgcttgctccgcg | XbaI | |
| CCT6-iF | cccaagcttgtgaggctggacgtgtacaa | HindIII | RNAi of CCT6 |
| CCT6-iR | cccaagcttgttatccaggatcccagccg | HindIII | |
| GAT2-iF | cccaagctttgggcatcggctacacaatt | HindIII | RNAi of GAT2 |
| GAT2-iR | cccaagcttcgcacgtcgatccatactca | HindIII | |
| ATG3-iF | gctctagaaatagctgccgaaaagacca | HindIII | RNAi of ATG3 or double RNAi of Rab2B and ATG3 |
| ATG3-iR | gctctagatgccatatgtgttgggaaga | HindIII | |
| Actin-iF | gctctagaatgtcggacgaggaacaaactg | XbaI | RNAi of actin |
| Actin-iR | gctctagacacagtatgtgtcacaccgtcac | XbaI | |
| $\alpha$-tubulin-iF | gctctagagcgttgaggatgatgcgttc | XbaI | RNAi of a-tubulin |
| $\alpha$-tubulin-iR | gctctagagtgagtgtgtcgagagcaca | XbaI | |
| $\beta$-tubulin-iF | gctctagaacctccgtaagttggctgtg | XbaI | RNAi of b-tubulin |

| | | | |
|---|---|---|---|
| β-tubulin-iR | gctctagatccatacctcgccagtgta | XbaI | |
| Raptor-iF | gctctagacgtgcacggtgttgtttcat | XbaI | RNAi of Raptor |
| Raptor-iR | gctctagactcattcgtaaaccgctgcg | XbaI | |
| mLST8-iF | gctctagagctctgctggagtgttgact | XbaI | RNAi of mLST8 |
| mLST8-iR | gctctagagagtcattcgtggcgattgc | XbaI | |
| CDC20-iF | gctctagagcattacacgccgattgcat | XbaI | RNAi of CDC20 |
| CDC20-iR | gctctagaccacctccccatcagttgac | XbaI | |
| H2A.Z-iF | gctctagaagcgaggaggtaaaactggc | XbaI | RNAi of H2A.Z |
| H2A.Z-iR | gctctagagctcttgtgcacaaatggca | XbaI | |
| ARP6-iF | gctctagaggttgcggtactacttgcct | XbaI | RNAi of ARP6 |
| ARP6-iR | gctctagaccggctttgcaccaatttca | XbaI | |
| RPA1-iF | gctctagaggatgaaaggggaagtggg | XbaI | RNAi of RPA1 |
| RPA1-iR | gctctagaaggggagcaataacagcacc | XbaI | |
| RPB1-iF | gctctagacgaaggagctgactcgatcc | XbaI | RNAi of RPB1 |
| RPB1iR | gctctagagacgtatcggagcggttgat | XbaI | |
| RPC1-iF | gctctagacctttccgcacgtttcgttt | XbaI | RNAi of RPC1 |
| RPC1-iR | gctctagatagccgtgcgaaaagtccat | XbaI | |
| Rab2B-F | gctctagaatgcagcagcaccattatgttttcaag | XbaI | Inducible OE of trypanosomal Rab2B |
| Rab2B-R | cgggatcctcagcagaagcagccactttcac | BamHI | |
| Rab2B-S21N-F | gtgggaaagaactgcctcctactg | / | Point mutagenesis to generate S21N mutation of Rab2B |
| Rab2B-S21N-R | cagtaggaggcagttctttcccac | / | |
| Rab2B-Q66L-F | gatacagctggactggaaagcttccg | / | Point mutagenesis to generate Q66L mutation of Rab2B |
| Rab2B-Q66L-R | cggaagctttccagtccagctgtatc | / | |
| Rab2B-bac-F | cgggatccatgcagcagcaccattatgttttcaag | BamHI | To express GST-tagged Rab2B in bacteria |
| Rab2B-bac-R | ggaattctcagcagaagcagccactttcac | EcoRI | |
| ActVHH-F | cccaagcttatggctcaggtgcagctggtg | HindIII | Expression of actin chromobody (nAc) and nuclear targeted nAc |
| ActVHH-NLS-F | cccaagcttatgcgcggccacaagcgctcgcgcgaggctc aggtgcagctggtg | HindIII | |

| | | | |
|---|---|---|---|
| ActVHH-R | cgggatccttacctgtacagctcgtccatgcc | BamHI | |
| P003-XbaI-F | gcaggtggagcaggttctagagtccaactggtggag | / | Point mutagenesis to create XbaI site in vector NLS-deGradFP |
| P003-XbaI-R | ctccaccagttggactctagaacctgctccacctgc | / | |
| P004-XbaI-F | gcaggtggagcaggttctagagtccaactggtggag | / | Point mutagenesis to create XbaI site in vector deGradFP |
| P004-XbaI-R | ctccaccagttggactctagaacctgctccacctgc | / | |
| nAc-deg--F | gctctagaatggctcaggtgcagctggtg | XbaI | Create vectors degrAdCT and NLS-degrAdCT |
| nAc-deg--R | cgggatcc<u>tta</u>tgaggagacggtgacctgg | BamHI | |
| nAc-bac-F | ggaattcatggctcaggtgcagctggtg | EcoRI | To express GST-tagged nAc in bacteria |
| nAc-bac-R | ccgctcgagttatgaggagacggtgacctgg | XhoI | |
| HsRab2A-F | cgggatccatggcgtacgcctatctcttcaag | BamHI | Transient express of HsRab2A |
| HsRab2A-R | ccgctcgagtcaacagcagccgcccc | XhoI | |
| HsRab2ASN-F | cacaggtgttggtaaaaactgcttattgctacag | / | Point mutagenesis to create S20N mutant of HsRab2A |
| HsRab2ASN-R | ctgtagcaataagcagttttaccaacacctgtg | / | |
| HsRab2AQL-F | tgggatacggcagggctcgaatcctttcgttcc | / | Point mutagenesis to create Q65L mutant of HsRab2A |
| HsRab2AQL-R | ggaacgaaaggattcgagccctgccgtatccca | / | |
| HsRab2B-F | cgggatccatgacttatgcttatctcttcaagtatatcatcatcg | BamHI | Transient expression of HsRab2B |
| HsRab2B-R | ataagaatgcggccgctcagcagcagccagagttg | NotI | |
| HsRab2BSN-F | acaggtgtggggaagaactgtctcctcctgcag | / | Point mutagenesis to create S20N mutant of HsRab2B |
| HsRab2BSN-R | ctgcaggaggagacagttcttccccacacctgt | / | |
| HsRab2BQL-F | tgggatacggctgggctcgaatccttccgttct | / | Point mutagenesis to create Q65L mutant of HsRab2B |
| HsRab2BQL-R | agaacggaaggattcgagcccagccgtatccca | / | |
